# Multi-ancestry GWAS of the electrocardiographic PR interval identifies 210 loci underlying cardiac conduction

**DOI:** 10.1101/712398

**Authors:** Ioanna Ntalla, Lu-Chen Weng, James H. Cartwright, Amelia Weber Hall, Gardar Sveinbjornsson, Nathan R. Tucker, Seung Hoan Choi, Mark D. Chaffin, Carolina Roselli, Michael R. Barnes, Borbala Mifsud, Helen R. Warren, Caroline Hayward, Jonathan Marten, James J. Cranley, Maria Pina Concas, Paolo Gasparini, Thibaud Boutin, Ivana Kolcic, Ozren Polasek, Igor Rudan, Nathalia M. Araujo, Maria Fernanda Lima-Costa, Antonio Luiz P. Ribeiro, Renan P. Souza, Eduardo Tarazona-Santos, Vilmantas Giedraitis, Erik Ingelsson, Anubha Mahajan, Andrew P. Morris, Greco M. Fabiola Del, Luisa Foco, Martin Gögele, Andrew A. Hicks, James P. Cook, Lars Lind, Cecilia M. Lindgren, Johan Sundström, Christopher P. Nelson, Muhammad B. Riaz, Nilesh J. Samani, Gianfranco Sinagra, Sheila Ulivi, Mika Kähönen, Pashupati P. Mishra, Nina Mononen, Kjell Nikus, Mark J. Caulfield, Anna Dominiczak, Sandosh Padmanabhan, May E. Montasser, Jeff R. O’Connell, Kathleen Ryan, Alan R. Shuldiner, Stefanie Aeschbacher, David Conen, Lorenz Risch, Sébastien Thériault, Nina Hutri-Kähönen, Terho Lehtimäki, Leo-Pekka Lyytikäinen, Olli T. Raitakari, Catriona L. K. Barnes, Harry Campbell, Peter K. Joshi, James F. Wilson, Aaron Isaacs, Jan A. Kors, Cornelia M. van Duijn, Paul L. Huang, Vilmundur Gudnason, Tamara B. Harris, Lenore J. Launer, Albert V. Smith, Erwin P. Bottinger, Ruth J. F. Loos, Girish N. Nadkarni, Michael H. Preuss, Adolfo Correa, Hao Mei, James Wilson, Thomas Meitinger, Martina Müller-Nurasyid, Annette Peters, Melanie Waldenberger, Massimo Mangino, Timothy D. Spector, Michiel Rienstra, Yordi J. van de Vegte, Pim van der Harst, Niek Verweij, Stefan Kääb, Katharina Schramm, Moritz F. Sinner, Konstantin Strauch, Michael J. Cutler, Diane Fatkin, Barry London, Morten Olesen, Dan M. Roden, M. Benjamin Shoemaker, J. Gustav Smith, Mary L. Biggs, Joshua C. Bis, Jennifer A. Brody, Bruce M. Psaty, Ken Rice, Nona Sotoodehnia, Alessandro De Grandi, Christian Fuchsberger, Cristian Pattaro, Peter P. Pramstaller, Ian Ford, J. Wouter Jukema, Peter W. Macfarlane, Stella Trompet, Marcus Dörr, Stephan B. Felix, Uwe Völker, Stefan Weiss, Aki S. Havulinna, Antti Jula, Katri Sääksjärvi, Veikko Salomaa, Xiuqing Guo, Susan R. Heckbert, Henry J. Lin, Jerome I. Rotter, Kent D. Taylor, Jie Yao, Renée de Mutsert, Arie C. Maan, Dennis O. Mook-Kanamori, Raymond Noordam, Francesco Cucca, Jun Ding, Edward G. Lakatta, Yong Qian, Kirill V. Tarasov, Daniel Levy, Honghuang Lin, Christopher H. Newton-Cheh, Kathryn L. Lunetta, Alison D. Murray, David J. Porteous, Blair H. Smith, Bruno H. Stricker, André Uitterlinden, Marten E. van den Berg, Jeffrey Haessler, Rebecca D. Jackson, Charles Kooperberg, Ulrike Peters, Alexander P. Reiner, Eric A. Whitsel, Alvaro Alonso, Dan E. Arking, Eric Boerwinkle, Georg B. Ehret, Elsayed Z. Soliman, Christy L. Avery, Stephanie M. Gogarten, Kathleen F. Kerr, Cathy C. Laurie, Amanda A. Seyerle, Adrienne Stilp, Solmaz Assa, M. Abdullah Said, M. Yldau van der Ende, Pier D. Lambiase, Michele Orini, Julia Ramirez, Stefan Van Duijvenboden, David O. Arnar, Daniel F. Gudbjartsson, Hilma Holm, Patrick Sulem, Gudmar Thorleifsson, Rosa B. Thorolfsdottir, Unnur Thorsteinsdottir, Emelia J. Benjamin, Andrew Tinker, Kari Stefansson, Patrick T. Ellinor, Yalda Jamshidi, Steven A. Lubitz, Patricia B. Munroe

## Abstract

The electrocardiographic PR interval reflects atrioventricular conduction, and is associated with conduction abnormalities, pacemaker implantation, atrial fibrillation (AF), and cardiovascular mortality^1,2^. We performed multi-ancestry (N=293,051) and European only (N=271,570) genome-wide association (GWAS) meta-analyses for the PR interval, discovering 210 loci of which 149 are novel. Variants at all loci nearly doubled the percentage of heritability explained, from 33.5% to 62.6%. We observed enrichment for genes involved in cardiac muscle development/contraction and the cytoskeleton highlighting key regulation processes for atrioventricular conduction. Additionally, 19 novel loci harbour genes underlying inherited monogenic heart diseases suggesting the role of these genes in cardiovascular pathology in the general population. We showed that polygenic predisposition to PR interval duration is an endophenotype for cardiovascular disease risk, including distal conduction disease, AF, atrioventricular pre-excitation, non-ischemic cardiomyopathy, and coronary heart disease. These findings advance our understanding of the polygenic basis of cardiac conduction, and the genetic relationship between PR interval duration and cardiovascular disease.

## Main text

The electrocardiogram is among the most common clinical tests ordered to assess cardiac abnormalities. Reproducible waveforms indicating discrete electrophysiologic processes were described over 100 years ago, yet the biological underpinnings of conduction and repolarization remain incompletely defined. The electrocardiographic PR interval reflects conduction from the atria to ventricles, across specialised conduction tissues such as the atrioventricular node and the His-Purkinje system. Pathological variation in the PR interval may indicate heart block or pre-excitation, both of which can lead to sudden death^2^. The PR interval also serves as a risk factor for AF and cardiovascular mortality^1–3^. Prior genetic association studies have identified 64 PR interval loci^4–13^. To enhance our understanding of the genetic and biological mechanisms of atrioventricular conduction, we performed GWAS meta-analyses of autosomal and X chromosome variants imputed mainly with the 1000 Genomes Project reference panel^14^ using an additive model and increased sample size. Our primary meta-analysis included 293,051 individuals of European (92.6%), African (2.7%), Hispanic (4%), and Brazilian (<1%) ancestries from 40 studies (**Supplementary Tables 1-3**). We also performed ancestry-specific meta-analyses (**Fig. 1**).

**Figure 1.**
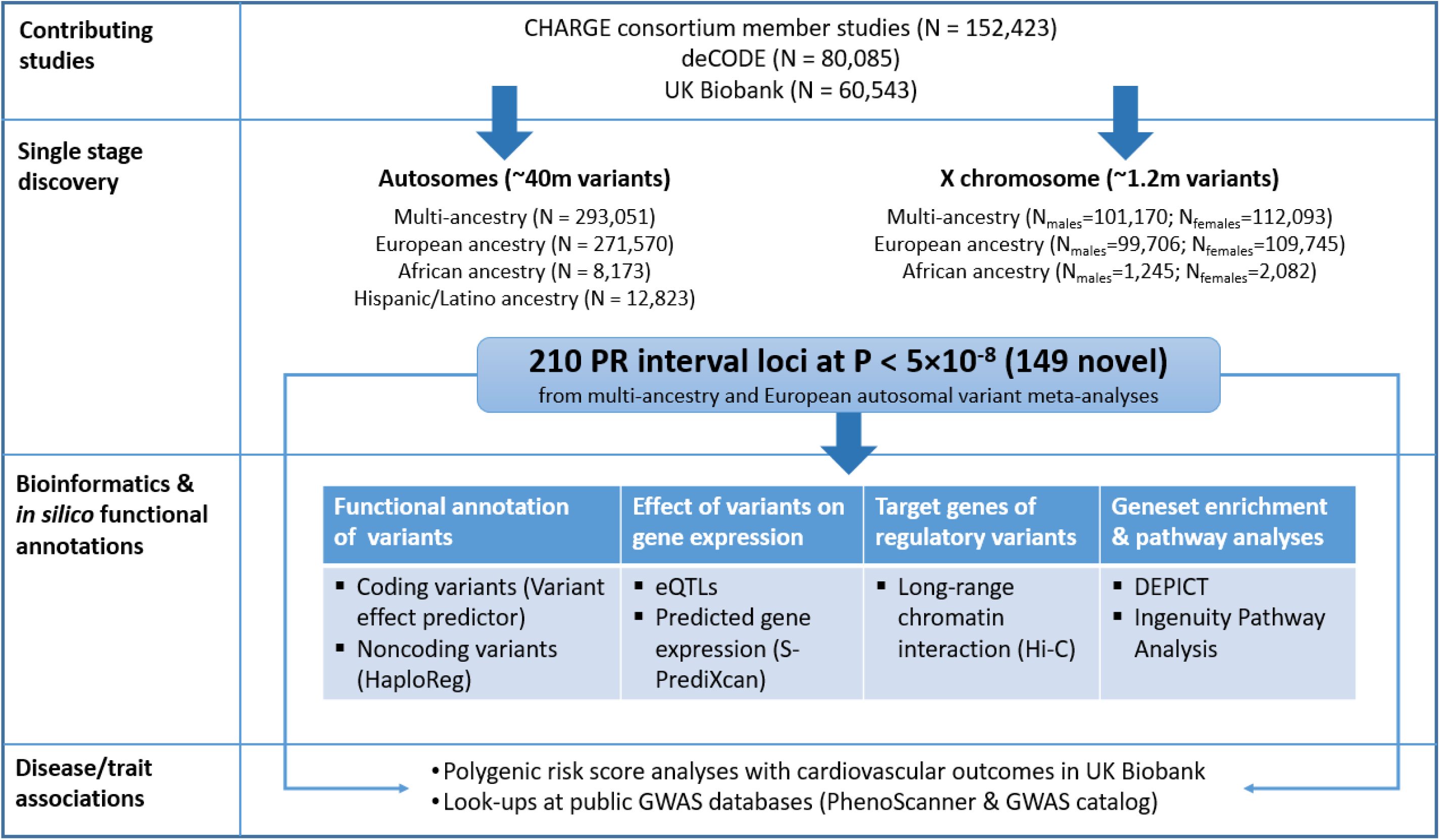
Overview of the study design. Figure includes overview of contributing studies, single-stage discovery approach, and downstream bioinformatics and *in silico* annotations we performed to link variants to genes, and polygenic risk score analysis to link variants to cardiovascular disease risk.

We identified a total of 210 genome-wide significant loci (P<5×10^−8^), of which 149 were not previously reported (**Table 1, Fig. 2**). Of the 149 novel loci, 141 were discovered in the multi-ancestry analysis, and 8 additional novel loci were identified in the European ancestry analysis (**Table 1, Fig. 2, Supplementary Tables 4-5, Supplementary Fig. 1-4**). We considered only variants present in >60% of the maximum sample size, a filtering criterion used to ensure robustness of associated loci (**Online Methods**). There was strong support for all 64 previously reported loci (61 at P<5×10^−8^ and 3 at P<1.1×10^−4^; **Supplementary Tables 6-7**). No additional novel loci were identified in African or Hispanic/Latino ancestry meta-analyses (**Supplementary Table 8, Supplementary Fig. 1 and 3**) or X chromosome meta-analyses (**Supplementary Fig. 5**). In secondary analyses, we examined the rank-based inverse normal transformed residuals of PR interval. Results of absolute and transformed trait meta-analyses were highly correlated (*ρ*>0.94, **Supplementary Tables 5, 9-10, Supplementary Fig. 6-7**).

**Table 1.**
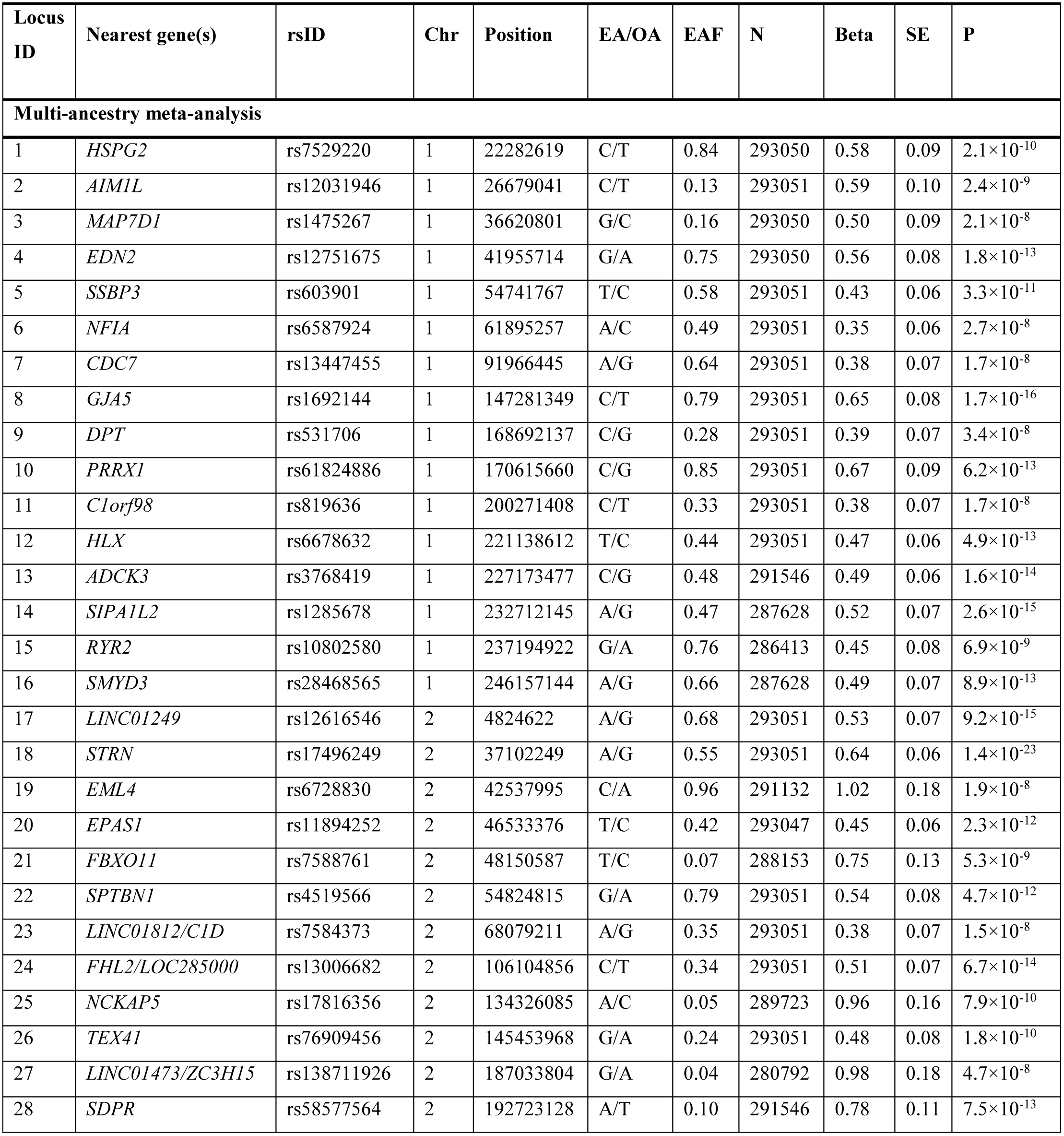
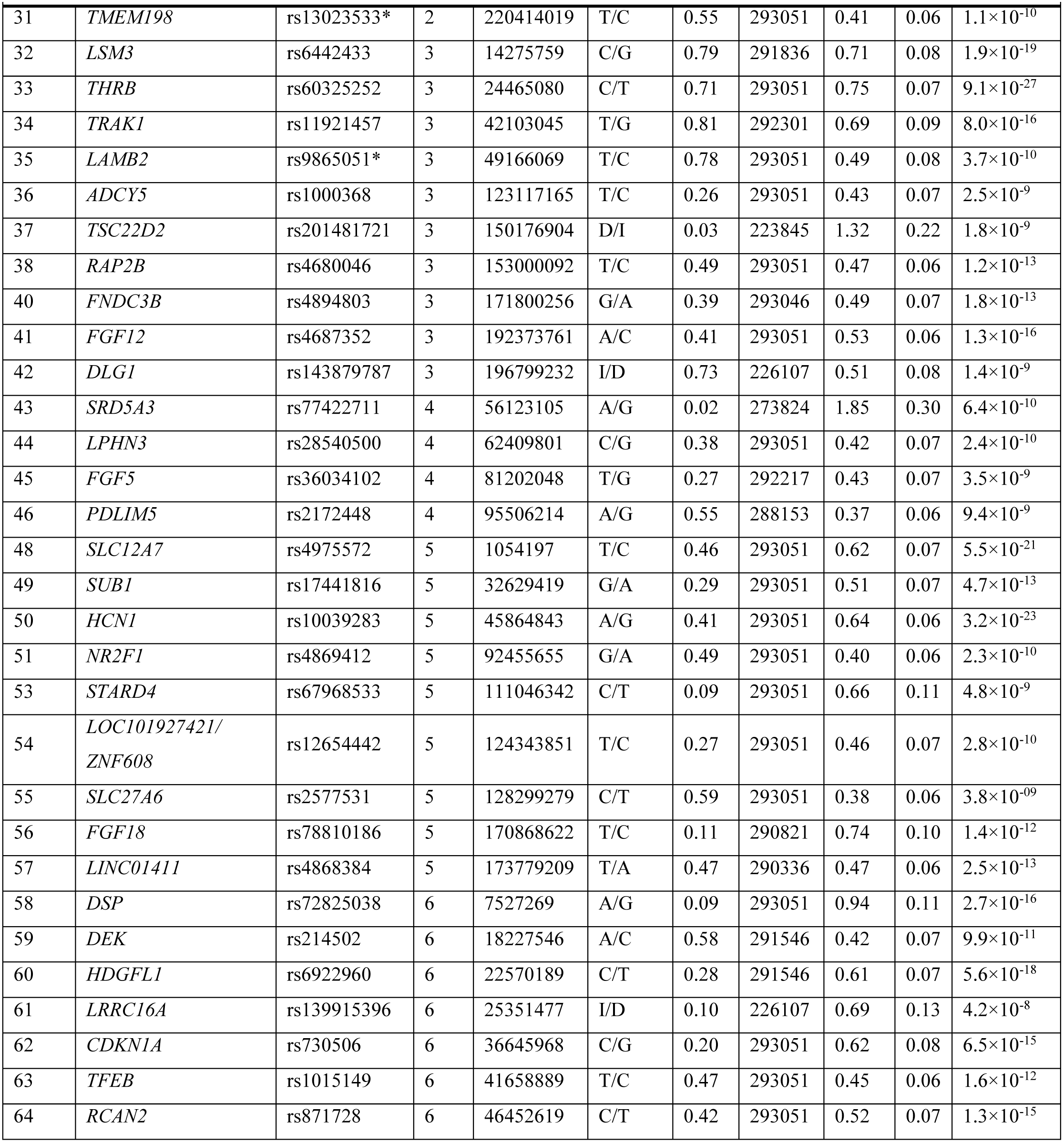
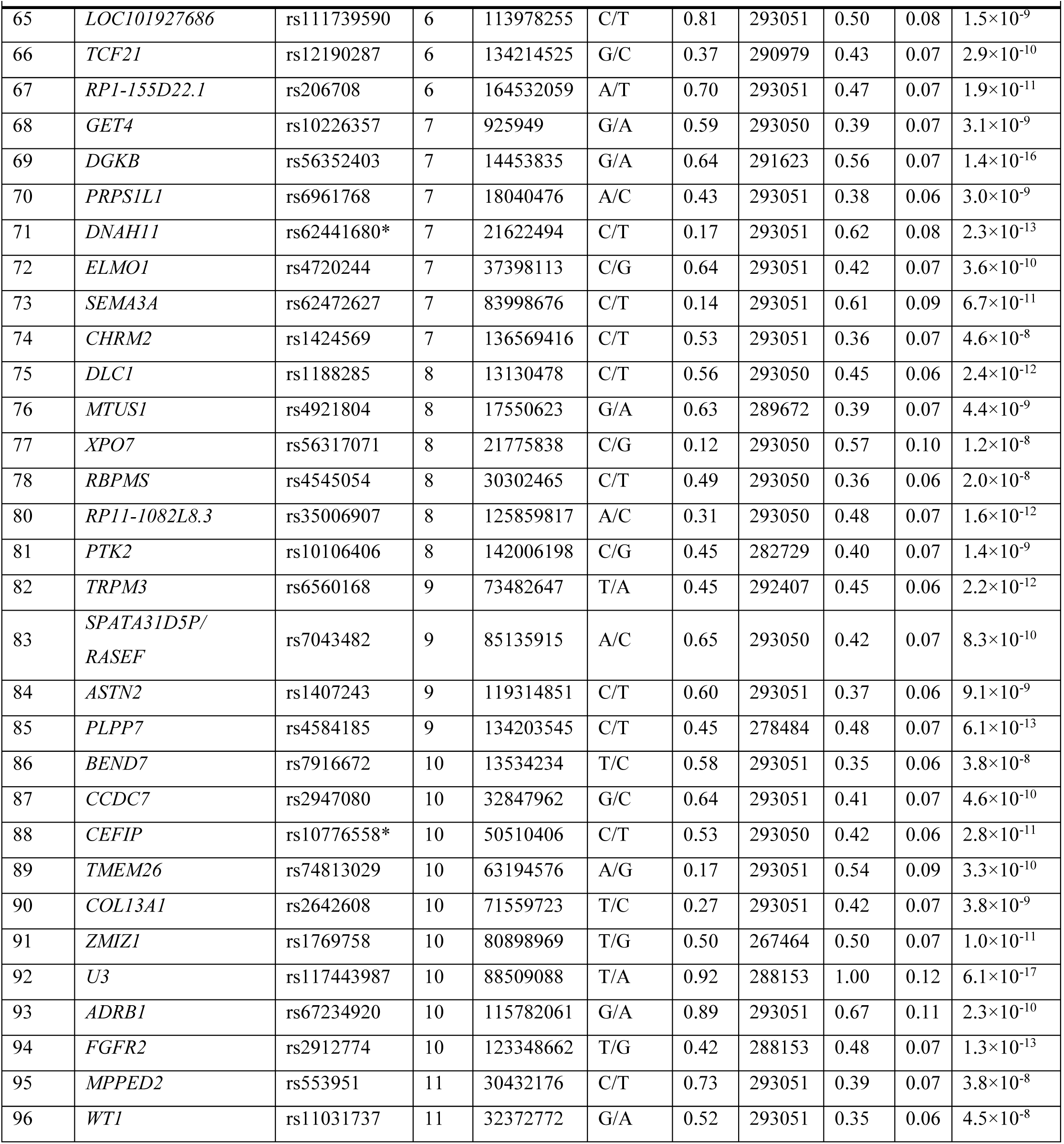
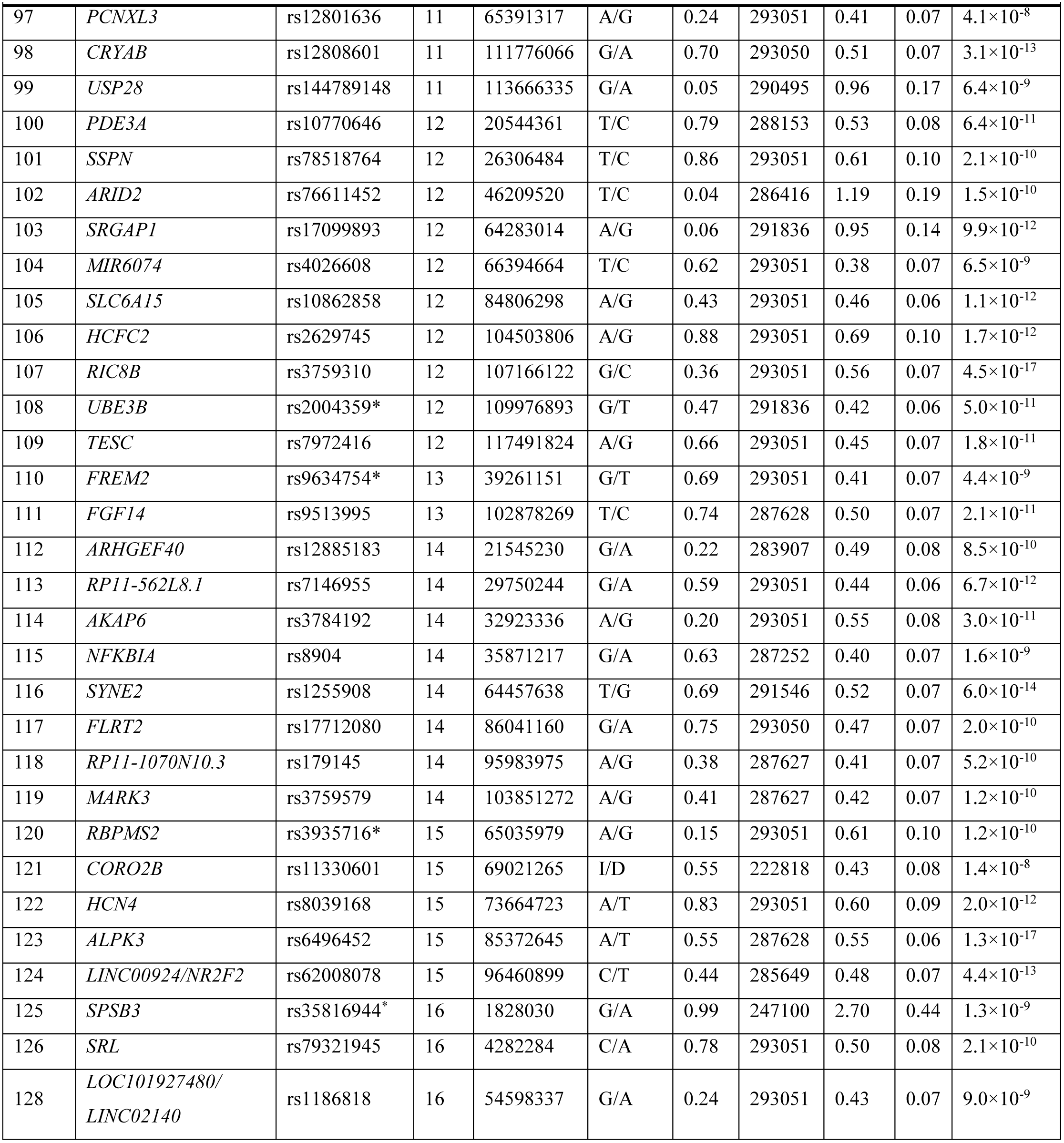
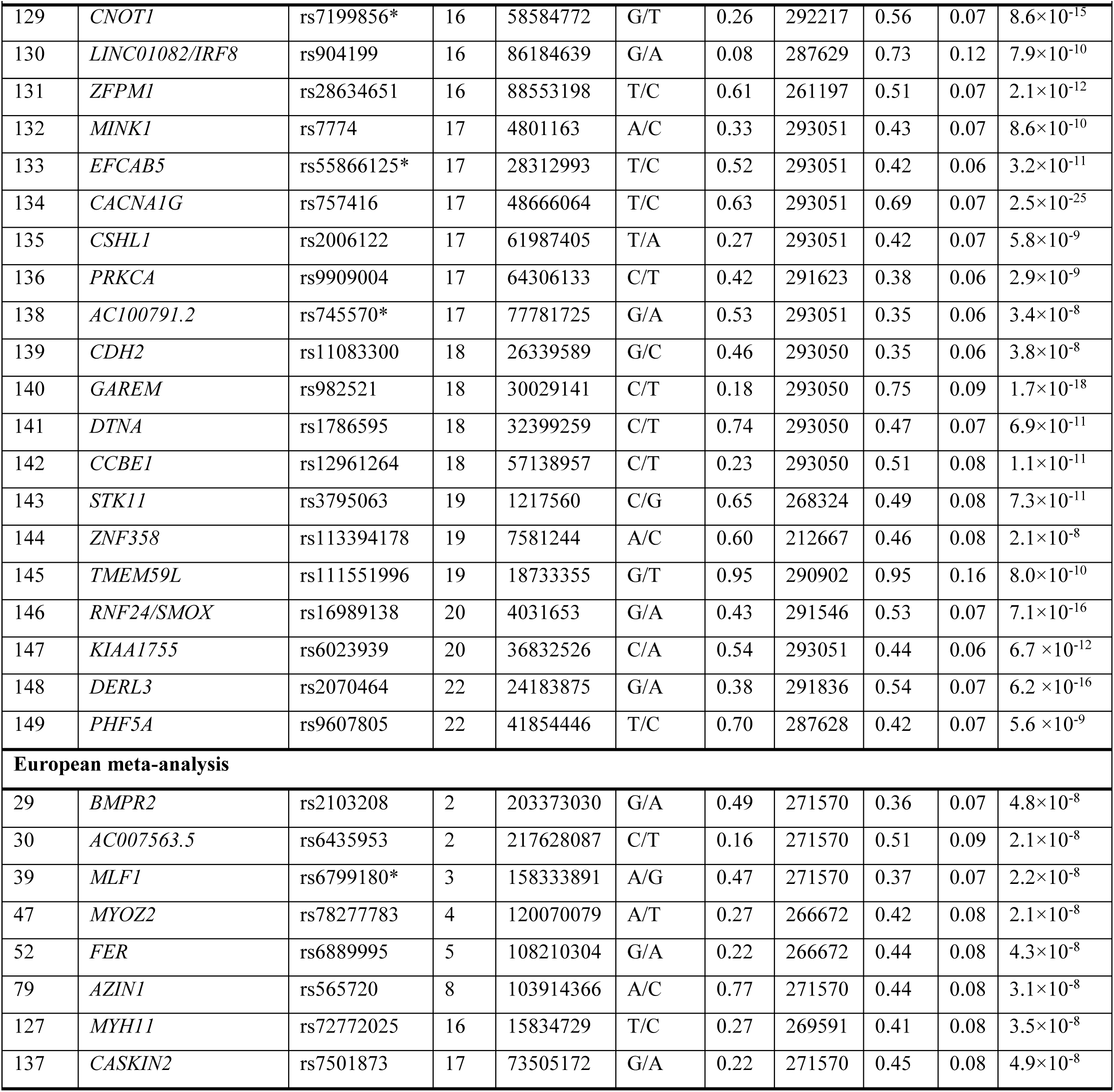
Novel genome-wide significant loci associated with PR interval (N = 149).

**Figure 2.**
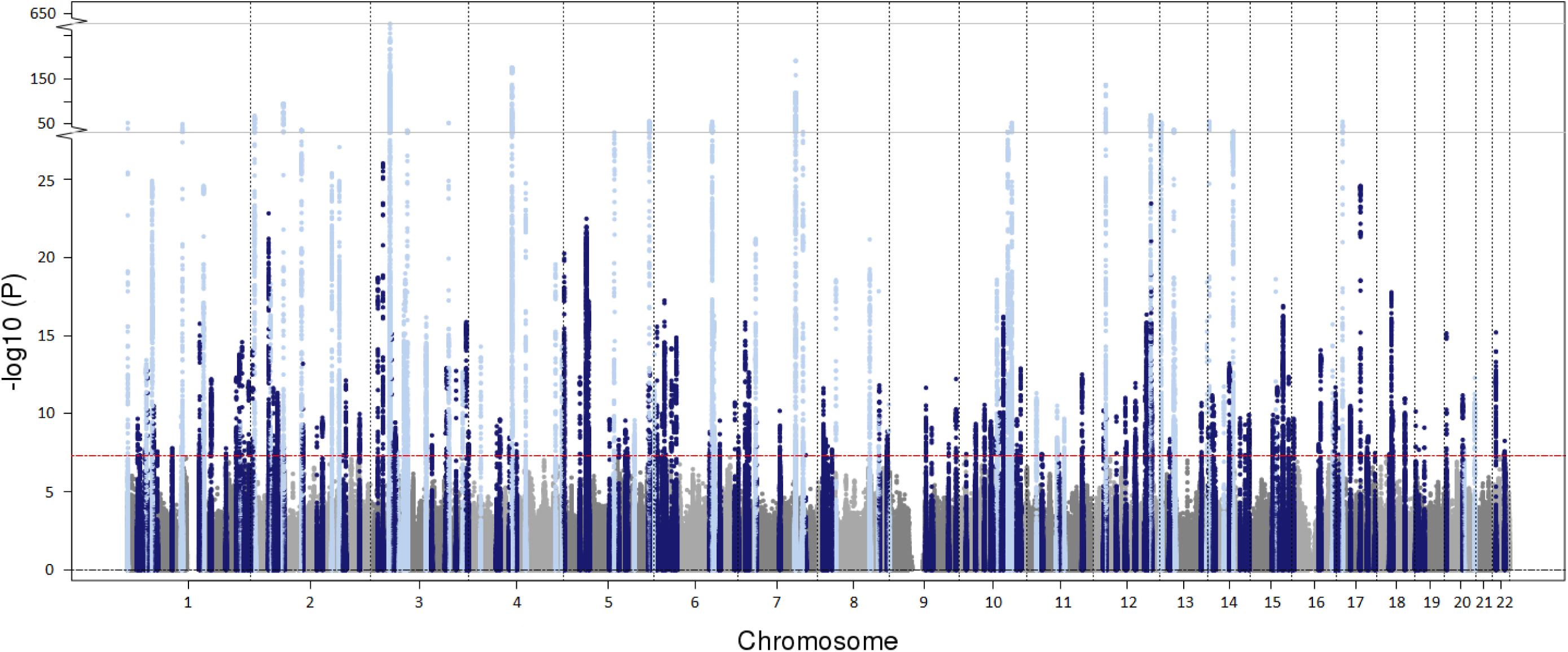
Manhattan plot of the multi-ancestry meta-analysis for PR interval. P values are plotted on the −log_10_ scale for all variants present in at least 60% of the maximum sample size. Associations of genome-wide significant (P < 5 × 10^−8^) variants at novel (N = 141) and previously reported loci (N = 61) are plotted in dark and light blue colours respectively.

**Figure 3.**
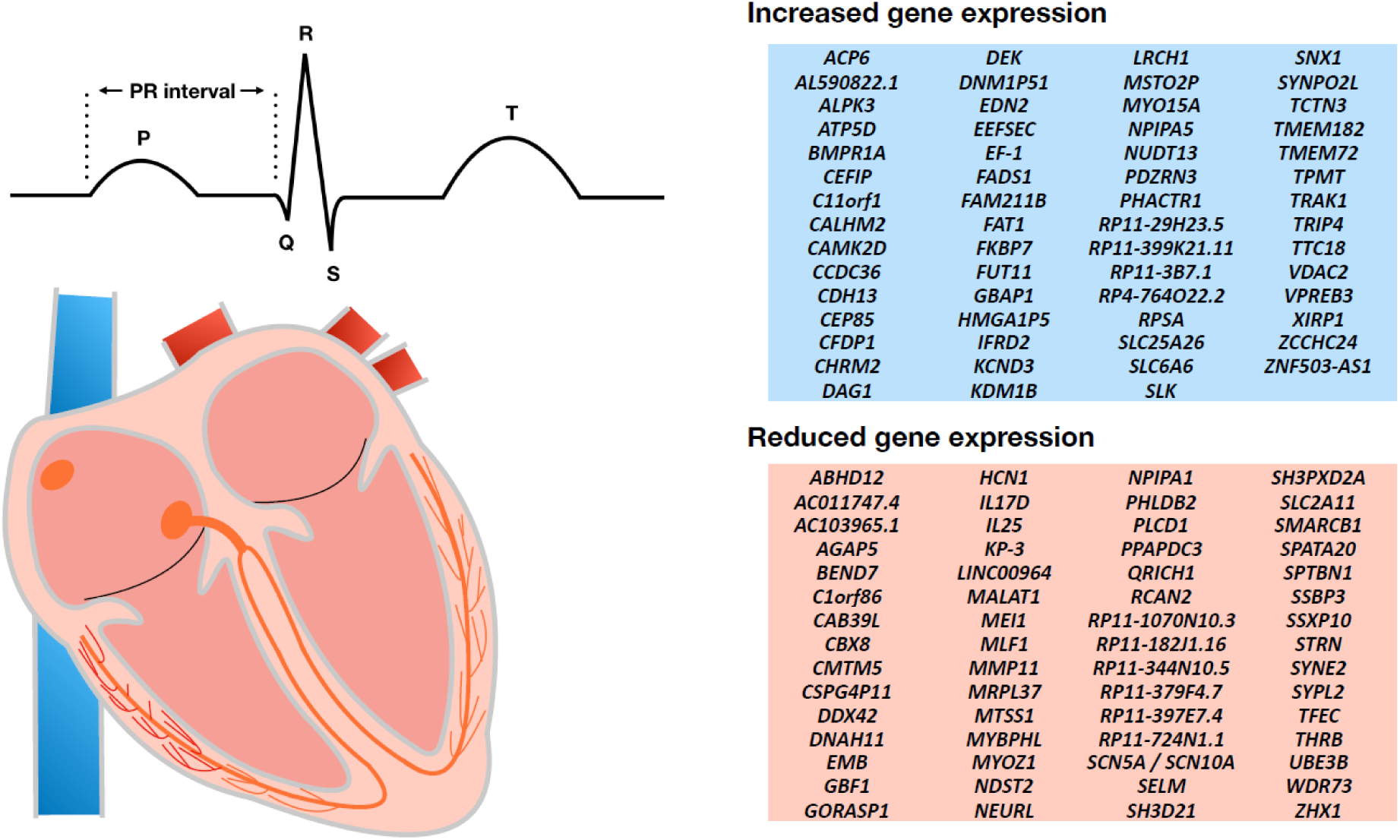
Plausible candidate genes of PR interval from S-PrediXcan. Diagram of standard electrocardiographic intervals and the heart. The electrocardiographic features are illustratively aligned with the corresponding cardiac conduction system structures (orange) reflected on the tracing. The PR interval (labeled) indicates conduction through the atria, atrioventricular node, His bundle, and Purkinje fibers. Right: The tables show 120 genes whose expression in the left ventricle (N=272) or right atrial appendage (N=264) was associated with PR interval duration in a transcriptome-wide analysis using S-PrediXcan and GTEx v7. Displayed genes include those with significant associations after Bonferroni correction for all tested genes at the two tissues with a P < 4.4×10^−6^ (=0.05/(5,977+5,366)). Longer PR intervals were associated with increased predicted expression of 59 genes (blue) and reduced expression of 61 genes (orange).

**Figure 4.**
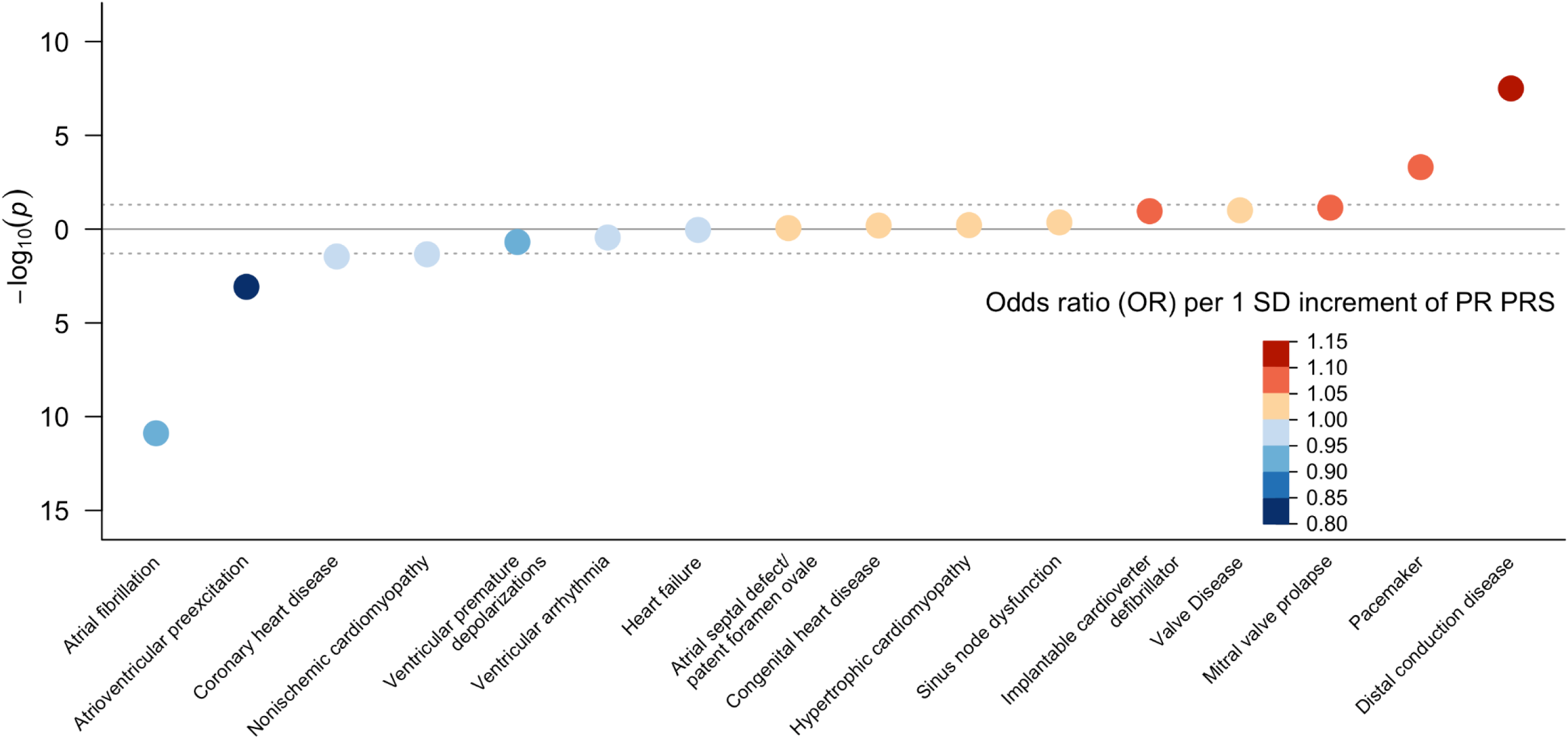
Bubble plot of phenome-wide association analysis of multi-ancestry PR interval polygenic risk score. Polygenic risk score was derived from the multi-ancestry meta-analysis results. Orange circles indicate that higher polygenic risk score of prolonged PR interval is associated with an increased risk of the condition, whereas blue circles indicate that higher score is associated with lower risks. The darkness of the colour reflects the effect size (odds ratio, OR) changes per 1 standard deviation increment of the polygenic risk score. Given correlation between traits, we did not establish a pre-specified significance threshold for the analysis and report nominal associations (P < 0.05).

**Figure 5.**
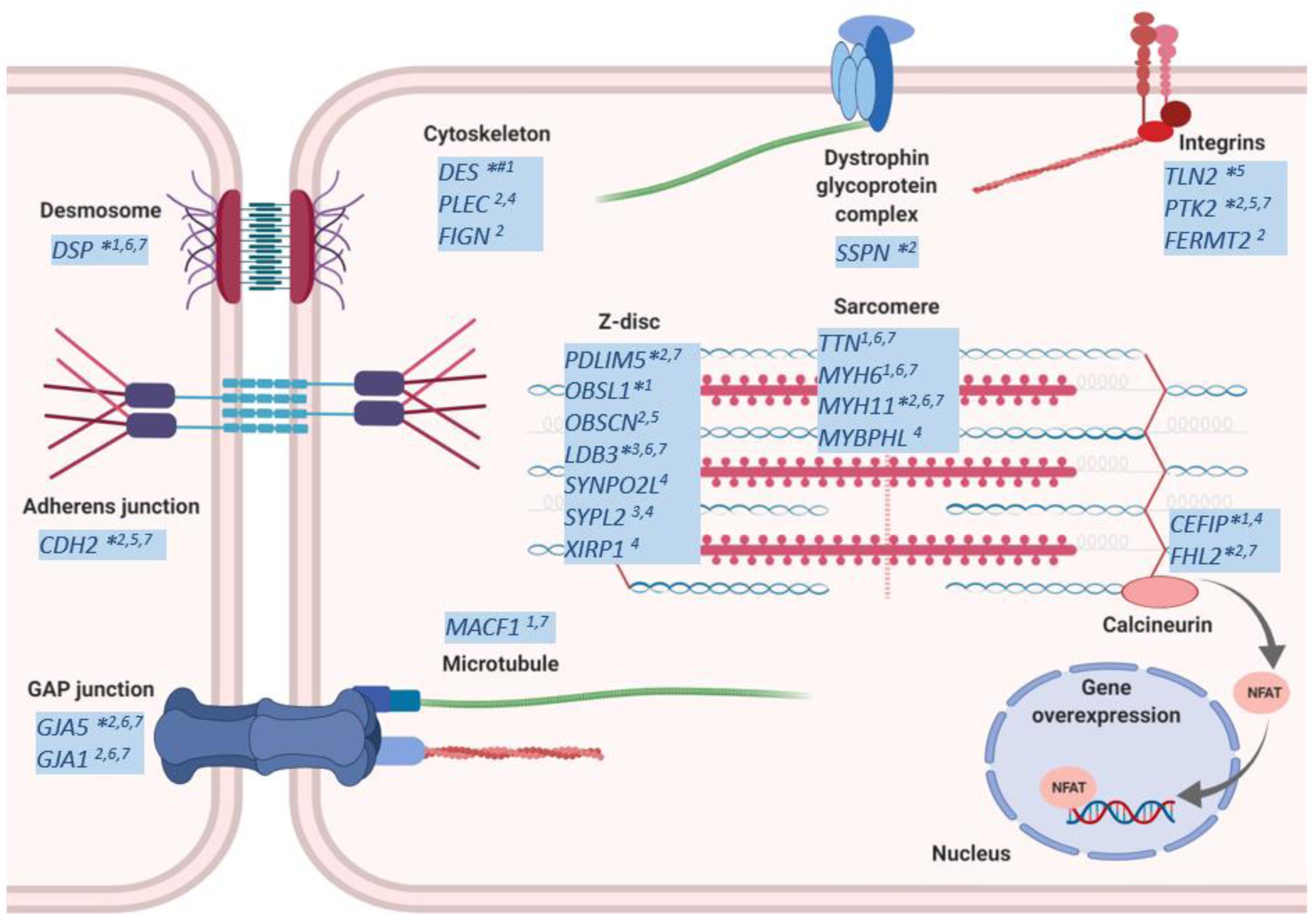
Candidate genes in PR interval loci encoding proteins involved in cardiac muscle cytoskeleton. Candidate genes or encoded proteins are indicated by a star symbol in the figure and listed in the table. More information about the genes is provided in Supplementary Tables 18-19. *Novel locus, # genome-wide significant locus in transformed trait meta-analysis. ^1^ Missense variant; ^2^ Nearest gene to the lead variant; ^3^ Gene within the region (r^2^>0.5); ^4^ Variant(s) in the locus are associated with gene expression in left ventricle and/or right atrial appendage; ^5^ Left ventricle best HiC locus interactor (RegulomeDB score ≤ 2); ^6^ Animal model; ^7^ Monogenic cardiovascular disease.

By applying joint and conditional analyses in the European meta-analysis data, we identified multiple independently associated variants (P_joint_<5×10^−8^ and r^2^<0.1) at 12 novel and 25 previously reported loci (**Supplementary Table 11**). The overall variant-based heritability (*h*^2^_g_) for the PR interval estimated in 59,097 unrelated European participants from the UK Biobank (UKB) with electrocardiograms was 18.2% (**Online Methods**). In the UKB, the proportion of *h*^2^_g_ explained by variation at all loci discovered in our analysis was 62.6%, compared to 33.5% when considering previously reported loci only.

The majority of the lead variants at the 149 novel loci were common (minor allele frequency, MAF>5%). We observed 6 low-frequency (MAF 1-5%) variants, and one rare (MAF<1%) predicted damaging missense variant (rs35816944, p.Ser171Leu) in *SPSB3* encoding SplA/Ryanodine Receptor Domain and SOCS Box-containing 3. SPSB3 is involved in degradation of SNAIL transcription factor, which regulates the epithelial-mesenchymal transition^15^, and has not been previously associated with cardiovascular traits. In total, we identified missense variants in genes at 12 novel and 6 previously reported loci (**Supplementary Table 12**). At *MYH6*, a previously described locus for PR interval^6,10^, sick sinus syndrome^16^, AF and other cardiovascular traits^17^, we observed a novel predicted damaging missense variant in *MYH6* (rs28711516, p.Gly56Arg). *MYH6* encodes the α-heavy chain subunit of cardiac myosin.

PR interval lead variants (or best proxy [r^2^>0.8]) at 39 novel and 23 previously reported loci were significant cis-eQTLs (at a 5% false discovery rate (FDR) in left ventricle (LV) and right atrial appendage (RAA) tissue samples from the Genotype-Tissue Expression (GTEx) project^18^ (**Supplementary Table 13)**. Variants at 21 novel loci were significant eQTLs in both tissues with consistent directionality of gene expression. We also performed a transcriptome-wide analysis to evaluate associations between predicted gene expression in LV and RAA with the PR interval. We identified 120 genes meeting our significance threshold (P<4.4×10^−6^, after Bonferroni correction); 26 genes were not localised at PR interval loci (≥500kb from a lead variant) representing potentially novel regions (**Supplementary Table 14, Supplementary Fig. 8**). Longer PR interval duration was associated with decreased levels of predicted gene expression for 61 genes, and increased levels for 59 genes (**Fig. 3**).

Most PR interval variants were annotated as non-coding. We therefore explored whether associated variants or proxies were located in transcriptionally active genomic regions. We observed enrichment for DNase I-hypersensitive sites in fetal heart tissue (P<9.36×10^−5^, **Supplementary Fig. 9**). Analysis of chromatin states indicated variants at 103 novel and 52 previously reported loci were located within regulatory elements that are present in heart tissues (**Supplementary Table 15**), providing support for gene regulatory mechanisms in specifying the PR interval. To identify distal candidate genes at PR interval loci, we assessed the same set of variants for chromatin interactions in a LV tissue Hi-C dataset^19^. Forty-eight target genes were identified (**Supplementary Table 16**). Variants at 38 novel loci were associated with other traits, including AF and coronary heart disease (**Supplementary Table 17, Supplementary Fig. 10**).

Candidate genes indicated by bioinformatics and *in silico* functional annotations at each novel locus are summarised in **Supplementary Tables 18-19,** and include 19 genes known to underlie monogenic cardiovascular diseases. Enrichment analysis of genes at PR interval loci using DEPICT^20^ indicated heart development (P=1.87×10^−15^) and actin cytoskeleton organisation (P=2.20×10^−15^) as the most significantly enriched processes (**Supplementary Table 20**). Ingenuity Pathway Analysis (IPA) supported heart development, ion channel signaling and cell-junction/cell-signaling amongst the most significant canonical pathways (**Supplementary Table 21**).

Finally, we evaluated associations between genetic predisposition to PR interval duration and 16 cardiac phenotypes chosen *a priori* using ∼309,000 unrelated UKB European participants not included in our meta-analyses^21^. We created a polygenic risk score (PRS) for PR interval using the multi-ancestry meta-analysis results (**Fig. 4**, **Supplementary Table 22**). Genetically determined PR interval prolongation was associated with higher risk of distal conduction disease (atrioventricular block; odds ratio [OR] per standard deviation 1.11, P=3.18×10^−8^) and pacemaker implantation (OR 1.06, P=0.0005). In contrast, genetically determined PR interval prolongation was associated with reduced risk of AF (OR 0.94, P=1.30×10^−11^) and atrioventricular pre-excitation (Wolff-Parkinson-White syndrome; OR 0.83, P=8.36×10^−4^). Genetically determined PR interval prolongation was marginally associated with a reduced risk of non-ischemic cardiomyopathy (OR=0.95, P=0.046) and coronary heart disease (OR 0.99, P=0.035). Results were similar when using a PRS derived using the European ancestry meta-analysis results (**Supplementary Fig. 11, Supplementary Table 22**).

To summarise, in meta-analyses of nearly 300,000 individuals we identified 210 loci, of which 149 were novel, underlying cardiac conduction as manifested by the electrocardiographic PR interval. Apart from confirming well-established associations in loci harbouring ion-channel genes, our findings further underscore the central importance of heart development and cytoskeletal components in atrioventricular conduction^10,12,13^. We also highlight the role of common variation at loci harboring genes underlying monogenic forms of heart disease in cardiac conduction.

We report signals in/near 13 candidate genes at novel loci with functional roles in cytoskeletal assembly (*DSP, DES, OBSL1, MYH11, PDLIM5, LDB3, FHL2, CEFIP, SSPN, TLN, PTK2, GJA5* and *CDH2*; **Fig. 5**). *DSP* and *DES* encode components of the cardiac desmosome, a complex involved in ionic communication between cardiomyocytes and maintenance of cellular integrity. Mutations in the desmosome are implicated in arrhythmogenic cardiomyopathy (ACM) and dilated cardiomyopathy (DCM)^22–26^. Conduction slowing is a major component of the pathophysiology of arrhythmia in ACM and other cardiomyopathies^27,28^. *OBSL1* encodes obscurin-like 1, which together with obscurin (OBSCN) is involved in sarcomerogenesis by bridging titin (TTN) and myomesin at the M-band^29^. *PDLIM5* encodes a scaffold protein that tethers protein kinases to the Z-disk, and has been associated with DCM in homozygous murine cardiac knockouts^30^. *FHL2* encodes calcineurin-binding protein four and a half LIM domains 2, which is involved in cardiac development by negatively regulating calcineurin/NFAT signaling in cardiomyocytes^31^. Missense mutations in *FHL2* have been associated with hypertrophic cardiomyopathy^32^. *CEFIP* encodes the cardiac-enriched FHL2-interacting protein located at the Z-disc, which interacts with *FHL2*. It is also involved in calcineurin–NFAT signaling, but its overexpression leads to cardiomyocyte hypertrophy^33^.

Common variants in/near genes associated with inherited arrhythmia syndromes were also observed, suggesting these genes also affect atrioventricular conduction and cardiovascular pathology in the general population. Apart from *DSP, DES, MYH11* and *GJA5* listed above, our analyses indicate 15 additional candidate genes (*ADRB1, ALPK3, BMPR1, BMPR2, CRYAB, DERL3, DNAH11, DTNA, ETV1, HCN4, MYOZ2, PDE3A, RYR2, SPEG, LDB3*) at novel loci causing Mendelian or other inherited forms of cardiovascular disease. Two genes we highlight are *HCN4* and *RYR2*. *HCN4* encodes a component of the hyperpolarization-activated cyclic nucleotide-gated potassium channel which specifies the sinoatrial pacemaker “funny” current, and is implicated in sinus node dysfunction, AF, and left ventricular noncompaction^34–36^. *RYR2* encodes a calcium channel component in the cardiac sarcoplasmic reticulum and is implicated in catecholaminergic polymorphic ventricular tachycardia^37^.

Genes with roles in autonomic signaling in the heart (*CHRM2, ADCY5*) were indicated from expression analyses. *CHRM2* encodes the M2 muscarinic cholinergic receptors that bind acetylcholine and are expressed in the heart^38^. Their stimulation results in inhibition of adenylate cyclase encoded by *ADCY5*, which in turn inhibits ion channel function. Ultimately, the signaling cascade can result in reduced levels of the pacemaker “funny” current in the sinoatrial and atrioventricular nodes, reduced L-type calcium current in all myocyte populations, and increased inwardly rectifying *I*_K.Ach_ potassium current in the conduction tissues and atria causing cardiomyocyte hyperpolarization^39^. Stimulation has also been reported to shorten atrial action potential duration and thereby facilitate re-entry, which may lead to AF^40–42^.

By constructing PRSs, we also observed that genetically determined PR interval duration is an endophenotype for several adult-onset complex cardiovascular diseases, the most significant of which are arrhythmias and conduction disorders. For example, our findings are consistent with previous epidemiologic data supporting a U-shaped relationship between PR interval duration and AF risk^1^. Although aggregate genetic predisposition to PR interval prolongation is associated with reduced AF risk, top PR interval prolonging alleles are associated with decreased AF risk (e.g., localized to the *SCN5A/SCN10A* locus) whereas others are associated with increased AF risk (e.g., localized to the *TTN* locus), consistent with prior reports^8^. These findings suggest that genetic determinants of the PR interval may identify distinct pathophysiologic mechanisms leading to AF, perhaps via specifying differences in tissue excitability, conduction velocity, or refractoriness. Future efforts are warranted to better understand the relations between genetically determined PR interval and specific arrhythmia mechanisms.

In conclusion, our study more than triples the reported number of PR interval loci, which collectively explain ∼62% of trait-related heritability. Our findings highlight important biological processes underlying atrioventricular conduction which include both ion channel function, and specification of cytoskeletal components. Our study also indicates that common variation in Mendelian cardiovascular disease genes contributes to population-based variation in the PR interval. Lastly, we observed that genetic determinants of the PR interval provide novel insights into the etiology of several complex cardiac diseases, including AF. Collectively, our results represent a major advance in understanding the polygenic nature of cardiac conduction, and the genetic relationship between PR interval duration and arrhythmias.

## Online Methods

### Contributing studies

A total of 40 studies (**Supplementary Note**) comprising 293,051 individuals of European (N=271,570), African (N=8,173), Hispanic (N=11,686), and Brazilian (N=485) ancestries contributed GWAS summary statistics for PR interval. All participating institutions and co-ordinating centres approved this project, and informed consent was obtained from all study participants. Study-specific design, sample quality control and descriptive statistics are provided in **Supplementary Tables 1-3**. For the majority of the studies imputation was performed for autosomal chromosomes and X chromosome using the 1000 Genomes (1000G) project^14^ reference panel or a most recently released haplotype version (**Supplementary Table 2**).

### PR interval phenotype and exclusions

The PR interval was measured in milliseconds from standard 12-lead electrocardiograms (ECGs), except in the UK-Biobank in which it was obtained from 4-lead ECGs (CAM-USB 6.5, Cardiosoft v6.51) recorded during a 15 second rest period prior to an exercise test (**Supplementary Note**). We excluded individuals with extreme PR interval values (<80ms or >320ms), second/third degree heart block, AF on the ECG, or a history of myocardial infarction or heart failure, Wolff-Parkinson-White syndrome, pacemakers, receiving class I and class III antiarrhythmic medications, digoxin, and pregnancy.

#### Study-level association analyses

We regressed the absolute PR interval on each genotype dosage using multiple linear regression with an additive genetic effect and adjusted for age, sex, height, body mass index, heart rate and any other study specific covariates. To account for relatedness, linear mixed effects models were used for family studies. To account for population structure, analyses were also adjusted for principal components of ancestry derived from genotyped variants after excluding related individuals. Analyses of autosomal variants were conducted separately for each ancestry group. X chromosome analyses were performed separately for males and females. Analyses using rank-based inverse normal transformed residuals of PR interval corrected for the aforementioned covariates were also conducted. Residuals were calculated separately by ancestral group for autosomal variants, and separately for males and females for X chromosome variants.

#### Centralized quality control

We performed quality control centrally for each result file using EasyQC version 11.4^43^. We removed variants that were monomorphic, had a minor allele count (MAC) <6, imputation quality metric <0.3 (imputed by MACH) or 0.4 (imputed by IMPUTE2), had invalid or mismatched alleles, were duplicated, or if they were allele frequency outliers (difference >0.2 from the allele frequency in 1000G project). We inspected PZ plots, effect allele frequency plots, effect size distributions, QQ plots, and compared effect sizes in each study to effect sizes from prior reports for established PR interval loci to identify genotype and study level anomalies. Variants with effective MAC (=2×N×MAF×imputation quality metric) <10 were omitted from each study prior to meta-analysis.

#### Meta-analyses

We aggregated summary level associations between genotypes and absolute PR interval from all individuals (N=293,051), and only from Europeans (N=271,570), African Americans (N=8,173), and Hispanic/Latinos (N=12,823) using a fixed-effects meta-analysis approach implemented in METAL (release on 2011/03/25)^44^. For the X chromosome, meta-analyses were conducted in a sex-stratified fashion. Genomic control was applied (if inflation factor λ_GC_>1) at the study level. Quantile–quantile (QQ) plots of observed versus expected –log_10_(P) did not show substantive inflation (**Supplementary Figs. 1-2**).

Given the large sample size we undertook a one-stage discovery study design. To ensure the robustness of this approach we considered for further investigation only variants reaching genome-wide significance (P<5×10^−8^) present in at least 60% of the maximum sample size (N_max_). We declared as novel any variants mapping outside the 64 loci previously reported (**Supplementary Note, Supplementary Table 6**). We grouped genome-wide significant variants into independent loci based on both distance (±500kb) and linkage disequilibrium (LD, r^2^<0.1) (**Supplementary Note**). We assessed heterogeneity in allelic effect sizes among studies contributing to the meta-analysis and among ancestral groups by the I^2^ inconsistency index^45^ for the lead variant in each novel locus. LocusZoom^46^ was used to create region plots of identified loci.

Meta-analyses (multi-ancestry [N=282,128], European only [N=271,570], and African [N=8,173]) of rank-based inverse normal transformed residuals of PR interval were also performed. Because not all studies contributed summary level association statistics of the transformed PR interval, we considered as primary the meta-analysis of absolute PR interval for which we achieved the maximum sample size. Any loci that met our significance criteria in the meta-analyses of transformed PR interval were not taken forward for downstream analyses.

#### Conditional and heritability analysis

Conditional and joint GWAS analyses were implemented in GCTA v1.91.3^47^ using summary level variant statistics from the European ancestry meta-analysis to identify independent association signals within PR interval loci. We used 59,097 unrelated (kinship coefficient >0.0884) UK Biobank participants of European ancestry as the reference sample to model patterns of LD between variants. We declared as conditionally independent any genome-wide significant variants in conditional analysis (*P*_joint_<5×10^−8^) not in LD (r^2^<0.1) with the lead variant in the locus.

Using the same set of individuals from UK Biobank, we estimated the aggregate genetic contributions to PR interval with restricted maximum likelihood as implemented in BOLT-REML^48^. We calculated the additive overall variant-heritability (*h*^2^_g_) based on 333,167 LD-pruned genotyped variants, as well as the *h*^2^_g_ of variants at PR interval associated loci only. Loci windows were based on both distance (±500kb) and LD (r^2^>0.1) around novel and previously reported variants (**Supplementary Note**). We then calculated the proportion of total *h*^2^_g_ explained at PR interval loci by dividing the *h*^2^_g_ estimate of PR interval loci by the total *h*^2^_g_.

#### Bioinformatics and in silico functional analyses

We use Variant Effect Predictor (VEP)^49^ to obtain functional characterization of variants including consequence, information on nearest genes and, where applicable, amino acid substitution and functional impact, based on SIFT^50^ and PolyPhen-2^51^ prediction tools. For non-coding variants, we assessed overlap with DNase I–hypersensitive sites (DHS) and chromatin states as determined by Roadmap Epigenomics Project ^52^ across all tissues and in cardiac tissues (E083, fetal heart; E095, LV; E104, right atrium; E105, right ventricle) using HaploReg v4^53^.

We assessed whether any PR interval variants were related to cardiac gene expression using GTEx^18^ version 7 cis-eQTL LV (N=272) and RAA (N=264) data. If the variant at a locus was not available in GTEx, we used proxy variants (r^2^>0.8). We report results only for associations at a false discovery rate (FDR) of 5%. We then evaluated the effects of predicted gene expression levels on PR interval duration using S-PrediXcan^54^. GTEx^18^ genotypes (variants with MAF>0.01) and normalized expression data in LV and RAA provided by the software developers were used as the training datasets for the prediction models. The prediction models between each gene-tissue pair were performed by Elastic-Net, and only significant (FDR 5%) models for prediction were included in our analysis. We used the European meta-analysis summary-level results (variants with at least 60% of maximum sample size) as the study dataset and then performed the S-PrediXcan calculator to estimate the expression-PR interval associations. In total, we tested 5,366 and 5,977 associations in LV and RAA, respectively. Significance threshold was set at P=4.4×10^−6^ (=0.05/(5,977+5,366)) to account for multiple testing corrections.

We applied GARFIELD (GWAS analysis of regulatory or functional information enrichment with LD correction)^55^ to analyse the enrichment patterns for functional annotations of the European meta-analysis summary statistics, using regulatory maps from the Encyclopedia of DNA Elements (ENCODE)^56^ and Roadmap Epigenomics^52^ projects. This method calculates odds ratios and enrichment P-values at different GWAS P-value thresholds (denoted T) for each annotation by using a logistic regression model accounting for LD, matched genotyping variants and local gene density with the application of logistic regression to derive statistical significance. Threshold for significant enrichment was set to P=9.36×10^−5^ (after multiple-testing correction for the number of effective annotations).

We identified potential target genes of regulatory variants using long-range chromatin interaction (Hi-C) data from the LV^19^. Hi-C data was corrected for genomic biases and distance using the Hi-C Pro and Fit-Hi-C pipelines according to Schmitt *et al.* (40kb resolution – correction applied to interactions with 50kb-5Mb span). We identified the promoter interactions for all potential regulatory variants in LD (r^2^>0.8) with our lead and conditionally independent PR interval variants and report the interactors with the variants with the highest regulatory potential (RegulomeDB≥2) to annotate the loci.

We performed a literature review, and queried the Online Mendelian Inheritance in Man (OMIM) and the International Mouse Phenotyping Consortium databases for all genes in regions defined by r^2^>0.5 from the lead variant at each novel locus. We further expanded the gene listing with any genes that were indicated by gene expression or chromatin interaction analyses. We performed look-ups for each lead variant or their proxies (r^2^>0.8) for associations (P<5×10^−8^) for common traits using both GWAS catalog^57^ and PhenoScanner v2^58^ databases. For AF, we supplemented the variant listing with a manually curated list of all overlapping variants (r^2^>0.7) with PR interval from two recently published GWASs^59,60^.

#### Gene set enrichment and pathway analyses

We used DEPICT (Data-driven Expression-Prioritized Integration for Complex Traits)^20^ to identify enriched pathways and tissues/cell types where genes from associated loci are highly expressed using all genome-wide significant (P<5×10^−8^) variants in our multi-ancestry meta-analysis present in at least 60% of N_max_ (N=20,076). To identify uncorrelated variants for PR interval, DEPICT performed LD-clumping (r^2^=0.1, window size=250kb) using LD estimates between variants from the 1000G reference data on individuals from all ancestries after excluding the major histocompatibility complex region on chromosome 6. Gene-set enrichment analysis was conducted based on 14,461 predefined reconstituted gene sets from various databases and data types, including Gene ontology, Kyoto encyclopedia of genes and genomes (KEGG), REACTOME, phenotypic gene sets derived from the Mouse genetics initiative, and protein molecular pathways derived from protein-protein interaction. Finally, tissue and cell type enrichment analysis was performed based on expression information in any of the 209 Medical Subject Heading (MeSH) annotations for the 37,427 human Affymetrix HGU133a2.0 platform microarray probes.

Ingenuity Pathway Analysis (IPA) was conducted using an extended list comprising 593 genes located in regions defined by r^2^>0.5 with the lead or conditionally independent variants for all PR interval loci, or the nearest gene. We further expanded this list by adding genes indicated by gene expression analyses. Only molecules and/or relationships for human or mouse or rat and experimentally verified results were considered. The significance P-value associated with enrichment of functional processes is calculated using the right-tailed Fisher’s exact test by considering the number of query molecules that participate in that function and the total number of molecules that are known to be associated with that function in the IPA.

#### Associations between genetically determined PR interval and cardiovascular conditions

We examined associations between genetic determinants of atrioventricular conduction and candidate cardiovascular diseases in unrelated individuals of European ancestry from UK Biobank (N∼309,000 not included in our GWAS meta-analyses) by creating PRSs for PR interval based on our GWAS results. We derived two PRSs. One was derived from the multi-ancestry meta-analysis results, and the other from the European meta-analysis results. We used the LD-clumping feature in PLINK v1.90^61^ (r^2^=0.1, window size=250kb, P=5×10^−8^) to select variants for each PRS. Referent LD structure was based on 1000G all ancestry, and European only data. In total, we selected 743 and 582 variants from multi-ancestry and European only meta-analysis results, respectively. We calculated the PRSs for PR interval by summing the dosage of PR interval prolonging alleles weighted by the corresponding effect size from the meta-analysis results. A total of 743 variants for the PRS derived from multi-ancestry results and 581 variants for the PRS derived from European results (among the variants with imputation quality >0.6) were included in our PRS calculations.

We selected candidate cardiovascular conditions *a priori*, which included various cardiac conduction and structural traits such as bradyarrhythmia, AF, atrioventricular pre-excitation, heart failure, cardiomyopathy, and congenital heart disease. We ascertained disease status based on data from baseline interviews, hospital diagnosis codes (ICD-9 and ICD-10), cause of death codes (ICD-10), and operation codes. Details of individual selections and disease definitions are described in **Supplementary Table 23**.

We tested the PRSs for association with cardiovascular conditions using logistic regression. We adjusted for enrolled age, sex, genotyping array, and phenotype-related principal components of ancestry. Given correlation between traits, we did not establish a pre-specified significance threshold for the analysis and report nominal associations (P<0.05).

There was no evidence of heterogeneity for any of the newly identified loci across individual studies (P_heterogeneity_ ≥ 0.001) or across ancestry groups (P_heterogeneity_ > 0.01).

Locus ID: unique locus identifier; Nearest gene(s): Nearest annotated gene(s) to the lead variant; rsID, variant accession number; Chr, chromosome; Position, physical position in build 37; EA, effect allele; OA, other allele; EAF, effect allele frequency; N, total sample size analyzed; beta, effect estimate is milliseconds; SE, standard error; P, P-value.

* Missense variant or variant in high LD (r^2^ > 0.8) with missense or splice site variant(s).

## URLs

1000 Genome Project: http://www.internationalgenome.org

BOLT-LMM: https://data.broadinstitute.org/alkesgroup/BOLT-LMM/

DEPICT: https://data.broadinstitute.org/mpg/depict/

DGIdb: http://www.dgidb.org

EasyQC: https://www.uni-regensburg.de/medizin/epidemiologie-praeventivmedizin/genetische-epidemiologie/software/#

FORGE: https://github.com/iandunham/Forge

GCTA: https://cnsgenomics.com/software/gcta/#Overview

GTEx: https://gtexportal.org/home/

HRC: http://www.haplotype-reference-consortium.org

IMPUTE2: http://mathgen.stats.ox.ac.uk/impute/impute_v2.html

Ingenuity Pathway Analysis software:

https://www.qiagenbioinformatics.com/products/ingenuity-pathway-analysis/

International Mouse Phenotyping Consortium: https://www.mousephenotype.org/

IPA: https://www.qiagenbioinformatics.com/products/ingenuity-pathway-analysis

LocusZoom: http://locuszoom.org/

MACH: http://csg.sph.umich.edu/abecasis/mach/tour/imputation.html

METAL: http://csg.sph.umich.edu/abecasis/metal/

OMIM: https://www.omim.org/

RegulomeDB: http://www.regulomedb.org

S-PrediXcan: https://github.com/hakyimlab/MetaXcan

UK Biobank: https://www.ukbiobank.ac.uk

## Author contributions

Interpreted results, writing and editing the manuscript: I.N., L.-C.W., S.A.L., and P.B.M. Conceptualisation and supervision of project: S.A.L. and P.B.M. Contributed to GWAS analysis plan: I.N., L.-C.W., H.R.W., Y.J., S.A.L., and P.B.M. Performed meta-analyses: I.N. and L.-C.W. Performed GCTA, heritability, geneset enrichment and pathway analyses, variant annotations: I.N. Performed polygenic risk score and gene expression analyses: S.H.C., M.D.C., and L.-C.W. Performed HiC analyses: I.N., M.R.B., B.M., and P.B.M. Performed gene literature review: I.N., L.-C.W., A.W.Hall, N.R.T., M.D.C., J.H.C., J.J.C., A.T., Y.J., S.A.L., and P.B.M. Contributed to study specific GWAS by providing phenotype, genotype and performing data analyses: J.M., I.R., C.H., P.G., M.Concas, T.B., O.P., I.K., E.T., N.M.A., R.P.S., M.F.L., A.L.P.R., A.M., V.Giedraitis, E.I., A.P.M., F.D.M., L.F., M.G., A.A.Hicks, J.P.C., L.Lind, C.M.L., J.Sundström, N.J.S., C.P.N., M.B.R., S.U., G.S., P.P.M., M.K., N.M., K.N., I.N., M.Caulfield, A.Dominiczak, S.P., M.E.M., J.R.O., A.R.S., K.Ryan, D.C., L.R., S.Aeschbacher, S.Thériault, T.L., O.T.R., N.H., L.Lyytikäinen, J.F.W., P.K.J., C.L.K.B., H.C., C.M.v., J.A.K., A.I., P.L.H., L.-C.W., S.A.L., P.T.E., T.B.H., L.J.L., A.V.S., V.Gudnason, E.P.B., R.J.F.L., G.N.N., M.H.P., A.C., H.M., J.W., M.Müller-Nurasyid, A.P., T.M., M.W., T.D.S., Y.J., M.Mangino, M.R., Y.J.V., P.H., N.V., K.Schramm, S.K., K.Strauch, M.F.S., B.L., C.R., D.F., M.J.C., M.Olesen, D.M.R., M.B.S., J.Smith, J.A.B., M.L.B., J.C.B., B.M.P., N.S., K.Rice, C.P., P.P.P., A.De Grandi, C.F., J.W.J., I.F., P.W.M., S.Trompet, S.W., M.D., S.B.F., U.V., A.S.Havulinna, A.J., K.Sääksjärvi, V.S., S.R.H., J.I.R., X.G., H.J.L., J.Y., K.D.T., R.N., R.d., D.O.M., A.C.M., F.C., J.D., E.G.L., Y.Q., K.V.T., E.J.B., D.L., H.L., C.H.N., K.L.L., A.D.M., D.J.P., B.H.Smith, B.H.Stricker, M.E.v, A.U., J.H., R.D.J., U.P., A.P.R., E.A.W., C.K., E.B., D.E.A., G.B.E., A.A., E.Z.S., C.L.A., S.M.G., K.F.K., C.C.L., A.A.S., A.S., S.Assa, M.A.S., M.Y.v., P.D.L., A.T., M.Orini, J.R., S.V.D., P.B.M., K.Stefansson, H.H., P.S., G.S., G.T., R.B.T., U.T., D.O.A., D.F.G. All authors read, revised and approved the manuscript.

