## Supplementary Note for "Multi-ancestry GWAS of the electrocardiographic PR interval identifies 210 loci underlying cardiac conduction"

1. **Description of participating studies**
2. **Supplementary Methods**

   **Definition of previously reported and novel loci**
3. **Supplementary Figures**

   **Supplementary Figure 1** Quantile-Quantile plots of autosomal variant results from meta-analyses of PR interval.

   **Supplementary Figure 2** Region plots of the 149 novel loci.

   **Supplementary Figure 3** Manhattan plot from the European (a), African (b), and Hispanic (c) meta-analyses of absolute PR interval.

   **Supplementary Figure 4** Correlation of association statistics for the lead variants at the 149 novel loci identified by the multi-ancestry and European ancestry meta-analyses of absolute PR interval.

   **Supplementary Figure 5** Quantile-Quantile plots of chromosome X variant results from meta-analyses of absolute PR interval.

**Supplementary Figure 6** Correlation of P-values across the meta-analyses of absolute and rank-based inverse normal transformed residuals of PR interval.

**Supplementary Figure 7** Genome-wide significant loci ($P = 5 \times 10^{-8}$) across meta-analyses of absolute and rank-based inverse normal transformed residuals PR interval.

**Supplementary Figure 8** Volcano plot of predicted gene expression analysis in cardiac tissues.

**Supplementary Figure 9** Enrichment of PR interval variants in DNAse I Hypersensitive sites.

**Supplementary Figure 10** Association of novel PR interval loci with other GWAS traits.

**Supplementary Figure 11** Bubble plot of phenome-wide association analysis of European-ancestry PR interval polygenic risk score.

4. **Study acknowledgments**

5. **Study funding**

6. **References**

### 1. Description of participating studies

Supplementary Table 1 indicates websites and references for further information on all contributing studies.

#### AMISH

The Old Order Amish (OOA) subjects included in this study were participants of several studies of cardiovascular health in relatively healthy volunteers from the OOA community of Lancaster County, PA and their family members. The studies were carried out at the University of Maryland as part of the Amish Complex Disease Research Program (ACDRP). The OOA population of Lancaster County, PA immigrated to the Colonies from Western Europe in the early 1700s. All study protocols were approved by the institutional review board at the University of Maryland and participating institutions. Informed consent was obtained from each of the study participants.

#### ARIC

The Atherosclerosis Risk in Communities (ARIC) Study[1] is a prospective community-based study of cardiovascular disease and its risk factors. At baseline (1987-89), 15,792 men and women age 45-64 were recruited from 4 communities in the US (Washington County, Maryland; Forsyth County, North Carolina; Jackson, Mississippi; Minneapolis suburbs, Minnesota). Participants were mostly white in the Minnesota and Washington County field centers, white and African American in Forsyth County, and exclusively African American in the Jackson field center. ECGs were recorded on MAC PC Personal Cardiographs (Marquette

Electronics Inc., Milwaukee, WI) and were subsequently submitted to a central reading center at the EPICORE Center (University of Alberta, Edmonton, Alberta, Canada) and thereafter to the Epidemiological Cardiology Research Center (EPICARE), Wake Forest University, Winston-Salem, NC. All ECGs were visually inspected for quality and legibility at their acquisition and by the reading centers and then stored in a digital format. The PR interval was determined as the mean duration in milliseconds from the P wave onset until the initiation of the QRS segment in the 12 ECG leads.

**BAMBUÍ**

A cohort study designed to identify predictors of adverse health events in the elderly. The study population comprises all residents of Bambuí (Minas Gerais, Brazil), aged 60 or more years (N=1,742). From these, 92.2% were interviewed and 85.9% underwent clinical examination, consisting of haematological and biochemical tests, serology for Trypanosoma cruzi, anthropometric and blood pressure measures and electrocardiogram. Cohort members undergo annual follow-up visits, which consist of an interview and verification of death certificates. Other procedures were repeated in selected years (2000, 2002 and 2008). From 1997 to 2007, during a mean follow-up of 8.6 years, 641 participants died and 96 (6.0%) were lost to follow-up.

**BioMe**

The Mount Sinai BioMe Biobank, founded in September 2007, is an ongoing, broadly consented EHR-linked bio- and data repository that enrolls participants non-selectively from the Mount Sinai Medical Center patient population. The BioMe Biobank draws from a population of over 70,000 inpatient and 800,000 outpatient visits annually from over 30 broadly selected clinical sites of the Mount Sinai Medical Center (MSMC). As of September 2017, BioMe has enrolled more than 42,000 patients that represent a broad racial, ethnic and socioeconomic diversity with a distinct and population-specific disease burden, characteristic of the communities served by Mount Sinai Hospital. BioMe participants are predominantly of African (AA, 24%), Hispanic/Latino (HL, 35%), European (EA, 32%), and other ancestry (OA, 10%). The BioMe Biobank Program operates under a Mount Sinai Institutional Review Board-approved research protocol. All study participants provided written informed consent.

**BRIGHT**

Participants of the BRIGHT Study were recruited from the Medical Research Council General Practice Framework and other primary care practices in the UK. Each case had a history of hypertension diagnosed prior to 60 years of age with confirmed blood pressure recordings corresponding to seated levels >150/100mmHg (1 reading) or mean of 3 readings >145/95 mmHg. The BRIGHT study focused on the recruitment of hypertensive individuals with BMI < 30. Sample selection for GWAS was based on DNA availability and quantity.

**Broad-AF**

The Broad AF Study is a collaborative project to investigate the genetic determinants of atrial fibrillation (AF), comprised of 17,517 AF cases and 10,987 referents from 26 studies. Details of study description, genotyping, and imputation were described previously[2]. Briefly, genetic variants were centrally genotyped on the Infinium PsychArray-24 v1.2 Bead Chip, and jointly called and quality controlled at the Broad Institute. After pre-imputation quality control, variants were imputed using the 1000 Genomes reference panel. A total of 3,461 individuals free of AF met the inclusion criteria of the current study and were included in the PR analysis. Individuals from the following participating studies were included: Vanderbilt University Medical Center Biobank (BioVU), Australian Familial AF Study, Danish AF Study, Groningen Genetics of Atrial Fibrillation (GGAF), Genetic Risk Assessment of Defibrillator Events Study (GRADE), Malmö Preventive Project (MPP-AF, and MPP-Echo), and Intermountain INSPIRE Registry. Study details were previously reported.[2]

**CAMP**

The MGH Cardiology and Metabolic Patient Cohort is comprised of 3850 subjects recruited from the ambulatory MGH Cardiology Practice between 2009 and 2012.

**CHRIS**

The Cooperative Health Research In South Tyrol (CHRIS) study is a population-based study with a longitudinal follow-up to investigate the genetic and molecular basis of age-related common chronic conditions and their interaction with life style and environment in the general population. The study was approved by the Ethics Committee of the Autonomous Province of Bolzano.

**CHS**

The Cardiovascular Health Study (CHS) is a population-based cohort study of risk factors for coronary heart disease and stroke in adults ≥65 years conducted across four field centers. The original predominantly European ancestry cohort of 5,201 persons was recruited in 1989-1990 from random samples of the Medicare eligibility lists; subsequently, an additional predominantly African-American cohort of 687 persons was enrolled for a total sample of 5,888.

Blood samples were drawn from all participants at their baseline examination and DNA was subsequently extracted from available samples. Genotyping was performed at the General Clinical Research Center's Phenotyping/Genotyping Laboratory at Cedars-Sinai among CHS participants who consented to genetic testing and had DNA available using the Illumina 370CNV BeadChip system (for European ancestry participants, in 2007) or the Illumina HumanOmni1-Quad_v1 BeadChip system (for African-American participants, in 2010).

CHS was approved by institutional review committees at each field center and individuals in the present analysis had available DNA and gave informed consent including consent to use of genetic information for the study of cardiovascular disease.

**CROATIA-Korcula**

The CROATIA-Korcula study sampled Croatians from the Adriatic island of Korcula, between the ages of 18 and 88. The fieldwork was performed in 2007 in the eastern part of the island, targeting healthy volunteers from the town of Korčula and the villages of Lumbarda, Žrnovo and Račišće.

**CROATIA-Split**

The CROATIA-Split study sampled Croatians from the city of Split, between the ages 18 and 85. The data was collected in 2008.

**deCODE**

The deCODE electrocardiogram (ECG) study was approved by the Data Protection Commission of Iceland and the National Bioethics Committee of Iceland (VSNb2015030024/03.01 with amendments). Written informed consent was obtained from individuals donating samples. Personal identifiers associated with medical information and samples were encrypted with a third-party encryption system as provided by the Data Protection Commission of Iceland. ECGs obtained in Landspitali - The National University Hospital of Iceland, Reykjavik, the largest and only tertiary care hospital in Iceland, have been digitally stored since 1998. For this analysis we used information on PR interval duration in milliseconds from individuals' first sinus rhythm ECG, including 80,085 individuals. We excluded individuals with permanent pacemakers or history of Wolff–Parkinson–White syndrome or atrioventricular block. We removed extreme outliers (PR-interval < 80 ms and PR-interval > 320ms) and adjusted the ECG measurements for sex, the R-R interval, year of birth and age at measurement. The genotypes in the deCODE study were derived from whole-genome sequencing of 8,383 Icelanders using Illumina standard TruSeq methodology to a mean depth of 30X, with subsequent imputation into 150,000 chip-typed individuals and their close relatives. The variants were then matched with variants found in 1000g phase 3 or the Haplotype Consortium reference panel. We tested the variants for association with PR interval duration in our data using linear regression. We assume that the quantitative trait follows a normal distribution with a mean that depends linearly on the expected allele at the SNP and a variance covariance matrix proportional to the kinship matrix. We used LD score regression to account for inflation in the test statistics.

**ERF**

The Erasmus Rucphen Family (ERF) study is comprised of a family-based cohort embedded in the Genetic Research in Isolated Populations (GRIP) program in the southwest of the Netherlands. The aim of this program is to identify genetic risk factors for the development of complex disorders. In ERF, twenty-two families that had a large number of children baptized in the community church between 1850 and 1900 were identified with the help of detailed genealogical records. All living descendants of these couples, and their spouses, were invited to take part in the study. Comprehensive interviews, questionnaires, and examinations were completed at a research center in the area; approximately 3,200 individuals participated. Examinations included 12 lead ECG measurements. Electrocardiograms were recorded on ACTA electrocardiographs (ESAOTE, Florence, Italy) and digital measurements of the PR interval were made using the Modular ECG Analysis System (MEANS). Data collection started in June 2002 and was completed in February 2005.

**FHS**

The objective of the Framingham Heart Study (FHS) was to identify the common factors or characteristics that contribute to CVD by following its development over a long period of time in a large group of participants. The researchers recruited 5,209 men and women between the ages of 30 and 62 years from the town of Framingham, Massachusetts, and began the first round of extensive physical examinations and lifestyle interviews. Since 1948, participants have continued to return to the study every two years for a detailed medical history, physical examination, and laboratory tests, and in 1971, the Study enrolled a second generation - 5,124 of the original participants' adult children and their spouses - to participate in similar examinations every four to eight years. In 1994, the need to establish a new study reflecting a more diverse community of Framingham was recognized, and the first Omni cohort of the Framingham Heart Study was enrolled. In April 2002 the Study entered a new phase, the enrollment of a third generation of participants, the grandchildren of the Original Cohort, who have been examined every four to eight years. In 2003, a second group of Omni participants was enrolled. Participants routinely received electrocardiograms at all research center visits. Electrocardiograms obtained at Original cohort Exam 11, Offspring cohort Exam 1, and Third Generation cohort Exam 1 were used in the PR analysis.

**GAPP**

GAPP is a population-based prospective cohort study involving a representative population-based sample of 2,170 healthy adults aged 25-41 years residing in the Principality of Liechtenstein. Exclusion criteria were the presence of cardiovascular disease, diabetes, obstructive sleep apnea and a body mass index >35kg/m2. A standardized 12-lead ECG was obtained in all participants. Details about the study have been described previously[3].

**FINCAVAS**

The purpose of the Finnish Cardiovascular Study (FINCAVAS) is to construct a risk profile - using genetic, haemodynamic and electrocardiographic (ECG) markers - of individuals at high risk of cardiovascular diseases, events and deaths. All patients scheduled for an exercise stress test at Tampere University Hospital and willing to participate have been recruited between October 2001 and December 2007. The final number of participants is 4,567. In addition to repeated measurement of heart rate and blood pressure, digital high-resolution ECG at 500 Hz was recorded continuously during the entire exercise test, including the resting and recovery phases. About 20% of the patients were examined with coronary angiography. Genetic variations known or suspected to alter cardiovascular function or pathophysiology were analysed to elucidate the effects and interactions of these candidate genes, exercise and commonly used cardiovascular medications.

**GRAPHIC**

Genetic Regulation of Arterial Pressure of Humans in the Community (GRAPHIC) Study: The GRAPHIC Study comprises 2024 individuals from 520 nuclear families recruited from the general population in Leicestershire, UK between 2003-2005 for the purpose of investigating the genetic determinants of blood pressure and related cardiovascular traits. Families were included if both parents aged 40-60 years and two offspring ≥18 years wished to participate. A detailed medical history was obtained from study subjects by standardized questionnaires and clinical examination was performed by research nurses following standard procedures. Measurements obtained included height, weight, waist-hip ratio, clinic and ambulatory blood pressure and a 12-lead ECG.

**GS-SFHS**

The GS:SFHS study recruited 23,960 participants aged 18–100 years between 2006–11; full details are reported elsewhere[4]. Participants came from across Scotland, with some family members from further afield. The sample was 59% female, with a wide range of ages and socio-demographic characteristics. Most (87%) participants were born in Scotland and 96% in the UK or Ireland. Mean family size (excluding 1400 singletons without any relations in the study) was 4.05 members; median was 3 (IQR 2–5). The largest family had 36 participating members, and participants were grouped in 5573 families. Genome-wide genotype data for nearly one million genetic variants has been measured on 10,000 selected participants. An important feature of GS:SFHS is the breadth and depth of phenotype information, including clinical and physical measures and detailed data on cognitive function, personality traits and mental health. Participants were asked a series of questions on smoking history, from which current smokers, former smokers and never smokers can be defined.

**HCHS/SOL**

The Hispanic Community Health Study/ Study of Latinos (HCHS/SOL) is a multicenter, community-based cohort study of U.S. Hispanics/Latinos. Goals of the study are to examine the prevalence of and risk factors for several disorders including heart, lung, blood, and kidney phenotypes. HCHS/SOL investigators sampled 16,415 males and females aged 18-74 years at baseline from four study communities: The Bronx, NY, Chicago, IL, Miami, FL, and San Diego, CA. HCHS/SOL recruitment centers were selected so that the study would include at least 2,000 participants in each of the following designations: Mexican, Puerto Rican, Dominican, Cuban, and Central and South American. 11,686 participants consented to genetic studies and are included in this analysis.

**Health 2000**

In Health 2000 (BRIF8901), a nationally representative sample of persons aged 30 or over was drawn from the nationwide population register in Finland. The survey focused on collecting information on health and functional capacity of the population, and included questionnaires, interviews and a comprehensive health examination.

**INGI-CARL**

INGI-CARL consisted of about 1000 subjects who were drawn from Carlantino, an isolated village of southern Italy. Ethics approval was obtained from the Ethics Committee of the "IRCCS Burlo Garofolo" in Trieste. Written informed consent was obtained from every participant of the study. The study population had undergone clinical and instrumental evaluations between 1998 and 2005. For all subjects, anthropometrics variables (such as height, weight, etc) were taken and a structured questionnaire about lifestyle and medical history was filled out. In addition, blood pressure, body-mass index, biochemical analyses, ECG and cardiovascular evaluation were collected.

**INGI-FVG**

The INGI-FVG cohort consisted of about 1700 subjects drawn from the project "Genetic Park of Friuli Venezia Giulia". This study examined 6 isolated villages in the North-east of Italy between 2008 and 2010. Ethics approval was obtained from the Ethics Committee of the "IRCCS Burlo Garofolo" in Trieste. Written informed consent was obtained from every participant of the study. The study population had undergone clinical and instrumental evaluations. For all subjects, anthropometrics variables (such as height, weight, etc) were taken and a structured questionnaire about lifestyle and medical history was filled out. In addition, blood pressure, body-mass index, biochemical analyses, ECG and cardiovascular evaluation were collected.

**JHS**

The JHS is a single-site cohort study of 5,306 extensively phenotyped African American women and men. Three clinical examinations have been completed, including the baseline examination, Examination 1 (2000–2004), Examination 2 (2005–2008), and Examination 3 (2009–2013), allowing comprehensive assessment of cardiovascular health and disease of the cohort at approximately four-year intervals. Ongoing monitoring of hospitalizations for cardiovascular events (coronary heart disease, heart failure and stroke) and deaths among cohort participants are accomplished by annual telephone follow-up interviews, surveillance of hospital discharge records (since 2000 for coronary heart disease and stroke, and since 2005 for heart failure), and vital records.

**KORA F3**

The KORA study is a series of independent population-based epidemiological surveys of participants living in the city of Augsburg, Southern Germany, or the two adjacent counties. All survey participants are residents of German nationality identified through the registration office and aged between 25 and 74 years at recruitment. The baseline survey KORA S3 was conducted in the years 1994/95. 3,006 participants from KORA S3 were reexamined in a 10-year follow-up (KORA F3) in the years 2004/05.

**KORA S4**

The KORA study is a series of independent population-based epidemiological surveys of participants living in the city of Augsburg, Southern Germany, or the two adjacent counties. All survey participants are residents of German nationality identified through the registration office and aged between 25 and 74 years at recruitment. The baseline survey KORA S4 was conducted in the years 1999-2001.

**LifeLines**

The LifeLines Cohort Study, and generation and management of GWAS genotype data for the LifeLines Cohort Study is supported by the Netherlands Organization of Scientific Research NWO (grant 175.010.2007.006), the Economic Structure Enhancing Fund (FES) of the Dutch government, the Ministry of Economic Affairs, the Ministry of Education, Culture and Science, the Ministry for Health, Welfare and Sports, the Northern Netherlands Collaboration of Provinces (SNN), the Province of Groningen, University Medical Center Groningen, the University of Groningen, Dutch Kidney Foundation and Dutch Diabetes Research Foundation.

**MESA**

The Multi-Ethnic Study of Atherosclerosis (MESA) is a study of the characteristics of subclinical cardiovascular disease (disease detected non-invasively before it has produced clinical signs and symptoms) and the risk factors that predict progression to clinically overt cardiovascular disease or progression of the subclinical disease. The cohort is a diverse, population-based sample of 6,814 asymptomatic men and women aged 45-84. Approximately

38 percent of the recruited participants are white, 28 percent African-American, 22 percent Hispanic, and 12 percent Asian (predominantly of Chinese descent). Participants were recruited during 2000-2002 from 6 field centers across the U.S. (at Wake Forest University; Columbia University; Johns Hopkins University; the University of Minnesota; Northwestern University; and the University of California – Los Angeles). All underwent anthropomorphic measurement and extensive evaluation by questionnaires at baseline, followed by 4 subsequent examinations at intervals of approximately 2-4 years. Age and sex were self-reported.

**MICROS**

The MICROS study is a population-based survey on adult volunteer participants who reside in three isolated villages in South Tyrol, Italy. These villages were selected because they had a small number of founders with old settlement, high rates of endogamy as well as slow/null population expansion. Extensive data was collected in 2002-03 regarding genealogy, and clinical measurements as well as collection of blood and urine samples and DNA isolation. An extensive standardized questionnaire was administered by interviewers to collect data on family history of disease and lifestyle exposures such as smoking and alcohol consumption. A serum sample was collected, prepared and stored at -80°C for subsequent analysis. The study was approved by the Ethics Committee of the Autonomous Province of Bolzano.

**NEO**

The NEO was designed for extensive phenotyping to investigate pathways that lead to obesity-related diseases. The NEO study is a population-based, prospective cohort study that includes 6,671 individuals aged 45–65 years, with an oversampling of individuals with overweight or obesity. At baseline, information on demography, lifestyle, and medical history have been collected by questionnaires. In addition, samples of 24-h urine, fasting and postprandial blood plasma and serum, and DNA were collected. Genotyping was performed using the Illumina HumanCoreExome chip, which was subsequently imputed to the 1000 genome reference panel. Participants underwent an extensive physical examination, including anthropometry, electrocardiography, spirometry, and measurement of the carotid artery intima-media thickness by ultrasonography. The collection of data started in September 2008 and completed at the end of September 2012. Participants are currently being followed for the incidence of obesity-related diseases and mortality.

**ORCADES**

The Orkney Complex Disease Study (ORCADES) is a family-based, cross-sectional study that seeks to identify genetic factors influencing cardiovascular and other disease risk in the isolated archipelago of the Orkney Isles in northern Scotland[5]. Genetic diversity in this population is decreased compared to Mainland Scotland, consistent with the high levels of endogamy historically. 2078 participants aged 16-100 years were recruited between 2005 and 2011, most having three or four grandparents from Orkney, the remainder with two Orcadian grandparents. Fasting blood samples were collected and many health-related phenotypes and environmental exposures were measured in each individual. All participants gave written informed consent and the study was approved by Research Ethics Committees in Orkney and Aberdeen (North of Scotland REC).

**PIVUS**

The Prospective Investigation of Vasculature in Uppsala Seniors study was initiated in 2001 to investigate the predictive power of different measurements of vascular characteristics for future cardiovascular events, and secondary aims included measurements of cardiac and metabolic function, as well as serum biomarkers and levels of environmental pollutants. All individuals aged 70 living in the community of Uppsala in Sweden were deemed eligible for the study. The subjects were selected from the community register and invited in randomized order between April 2001 and June 2004. They received an invitation letter for participation within 2 months of their 70th birthday. Of the 2,025 subjects invited, 1,016 (507 male, 509 female) subjects agreed to participate. The participants were asked to answer a questionnaire about their medical history, smoking habits and regular medication.

**PREVEND**

PREVEND (Prevention of Renal and Vascular End-stage Disease) study is an ongoing prospective study investigating the natural course of increased levels of urinary albumin excretion and its relation to renal and cardiovascular disease. Details of the protocol have been described elsewhere.

**PROSPER**

All data come from the PROspective Study of Pravastatin in the Elderly at Risk (PROSPER). A detailed description of the study has been published elsewhere. PROSPER was a prospective multicenter randomized placebo-controlled trial to assess whether treatment with pravastatin diminishes the risk of major vascular events in elderly. Between December 1997 and May 1999, we screened and enrolled subjects in Scotland (Glasgow), Ireland (Cork), and the Netherlands (Leiden). Men and women aged 70-82 years were recruited if they had pre-existing vascular disease or increased risk of such disease because of smoking, hypertension, or diabetes. A total number of 5,804 subjects were randomly assigned to pravastatin or placebo. A large number of prospective tests were performed including Biobank tests and cognitive function measurements. A whole genome wide screening has been performed in the sequential PHASE project. Of 5,763 subjects DNA was available for genotyping. Genotyping was performed with the Illumina 660K beadchip, after QC (call rate <95%) 5,244 subjects and 557,192 SNPs were left for analysis. These SNPs were imputed with IMPUTE software based on the 1000 Genomes Phase 3 panel. The study was approved by the institutional ethics review boards of centers of Cork University (Ireland), Glasgow University (Scotland) and Leiden University Medical Center (the Netherlands) and all participants gave written informed consent.

**RS**

Rotterdam Study (RS) is a prospective population-based cohort study. Details regarding design, objectives, and methods of the Rotterdam Study have been described in detail[6]. In short, the Rotterdam study started in 1989 with an initial cohort of 7,983 persons (out of 10,215 invitees; response rate 78%) 55 years of age or older living in the Ommoord district in the city of Rotterdam in the Netherlands. In 2000, 3,011 participants (out of 4,472 invitees, response rate 67%) who had become 55 years of age or moved into the study district were added to the cohort. Approximately every 4-5 years follow-up examinations are conducted. Examinations consist of a home interview and an extensive set of test at a research facility in the study district.

By linking the general practitioners' and municipality records to the study database, participants are continuously monitored for major morbidity and mortality.

**SardiNIA**

To identify genetic bases for prominent age-associated changes, including cardiovascular risk factors and determinants of personality traits, in a founder population. The results of the study will extend the studies of aging-associated conditions of outbred populations.

**SHIP/SHIP-Trend**

The Study of Health In Pomerania (SHIP) and SHIP-TREND both represent population-based studies. The Study of Health in Pomerania is a prospective longitudinal population-based cohort study in Western Pomerania assessing the prevalence and incidence of common diseases and their risk factors. Participants aged 20 to 79 with German citizenship and principal residency in the study area were recruited from a random sample of residents living in the three local cities, 12 towns as well as 17 randomly selected smaller towns. Individuals were randomly selected stratified by age and sex in proportion to population size of the city, town or small towns, respectively. A total of 4,308 participants were recruited between 1997 and 2001 in the SHIP cohort. Individuals were invited to the SHIP study centre for a computer-assisted personal interviews and extensive physical examinations.

**TwinsUK**

There are currently >13,500 twins registered participants in the TwinsUK study, of which over 9,000 are actively participating. The twins are aged 16 to 100 with approximately equal numbers of identical (MZ) and non-identical (DZ) twins and are predominantly female (80%) for historical reasons. Clinical, physiological, behavioural and lifestyle data is collected at either twin visits or via self-administered questionnaires, which volunteers complete either once or twice a year via the post or email. All studies have ethical approval from the Guy's and St Thomas' (GSTT) Ethics Committee.

**UK Biobank**

UK Biobank (UKB, www.ukbiobank.ac.uk) is a large longitudinal biobank study in the United Kingdom which was established to improve understanding of the genetic and environmental causes of common diseases including CVDs. In addition to self-reported disease outcomes and extensive health and life-style questionnaire data, UKB participants are being tracked through their NHS records and national registries (including cause of death and Hospital Episode Statistics). In 2017, UKB released the genotypes of 488,377 participants profiled with a custom SNP array. Genotyping QC was performed centrally by UKB, and genotypes imputed to Haplotype Reference Consortium (HRC) panel were released for 488,377 participants[7].

The PR interval was obtained from 4-lead ECGs (CAM-USB 6.5, Cardiosoft v6.51) recorded during a 15 second rest period prior to an exercise test while subjects were sitting on a stationary bike (eBike, Firmware v1.7). Electrodes were placed on the right and left antecubital fossae, and left and right wrist and the ECG was sampled at 500 Hz; lead I was used to derive the PR interval. To reduce the influence of noise, the PR interval was measured from a signal averaged ECG waveform computed from the heartbeats available in the 15s trace. For this, we first identified the QRS-complexes using fully automatic in-house algorithms [8,9]. Before averaging, we removed ectopic beats and artefacts, as well as beats with an RR interval longer or shorter than 10ms compared to the mean RR interval were not included for averaging. A signal-averaged heartbeat was then computed from the remaining beats provided that the number of available beats was not less than 5. Finally, the PR interval was automatically measured as the interval between the onset of the P-wave and the onset of the QRS complex from the signal-averaged heartbeat. Cases with arbitrary P- or QRS-waves were reviewed manually and corrected or removed if necessary.

**ULSAM**

The Uppsala Longitudinal Study of Adult Men (ULSAM) cohort is a study of healthy elderly men in the Uppsala region of Sweden. It was initiated as a health screen focused at identifying metabolic risk factors for cardiovascular disease. In 1970, all 50 year old men living in Uppsala were invited to participate. Of these, 82% of them participated initially. The cohort was subsequently invited back at ages 60, 70 and 77. This study included the collection of a wide range of phenotypes, including blood pressure, insulin metabolism, weight and height, lipid markers, diet, cognitive function and socio-economic factors.

**WHI**

The Women's Health Initiative (WHI) is a long-term national health study focused on strategies for preventing heart disease, breast and colorectal cancer, and osteoporotic fractures in postmenopausal women. Launched in 1993, the WHI enrolled 68,132 women aged 50-79 into one or more randomized Clinical Trials (CT), testing the health effects of hormone therapy (HT), dietary modification (DM), and/or calcium and Vitamin D supplementation (CaD). The genetic data used in this paper was generated by six ancillary studies: GWAS of Hormone Treatment and CVD and Metabolic Outcomes in WHI: Genomics and Randomized Trials Network (GARNET), Genome-wide Association Study of Nonsynonymous SNPs in Colon Cancer: Genetics and Epidemiology of Colorectal Cancer Consortium (GECCO), Genome-wide Association Study to Identify Genetic Components of Hip Fracture (HIPFX), Long Life Study (LLS), Modification of Particulate Matter-Mediated Arrhythmogenesis in Populations (MOPMAP), and Women's Health Initiative Memory Study (WHIMS). GARNET[10] is a case-control study of coronary heart disease, stroke, venous thromboembolism, and incident diabetes. GECCO[11] and HIPFX[12] are case-control studies of colon cancer and hip fracture. LLS[13] is a longitudinal study of the WHI Extension II Medical Records Cohort. MOPMAP[14] is a case-control study of ventricular ectopy. WHIMS[15] is a longitudinal study of cognitive decline, mild cognitive impairment, and dementia of the HT cohort.

**YFS**

The Young Finns study (YFS) is a population-based follow up-study started in 1980. The main aim of the YFS is to determine the contribution made by childhood lifestyle, biological and psychological measures to the risk of cardiovascular diseases in adulthood. In 1980, over 3,500 children and adolescents all around Finland participated in the baseline study. The follow-up studies have been conducted mainly with 3-year intervals. The 27-year follow-up study was conducted in 2007 (ages 30-45 years) with 2,204 participants. The study was approved by the local ethics committees (University Hospitals of Helsinki, Turku, Tampere, Kuopio and Oulu) and was conducted following the guidelines of the Declaration of Helsinki. All participants gave their written informed consent.

### 2. Supplementary Methods - Definition of previously reported and novel loci

A total of 98 uncorrelated ($r^2 < 0.1$) variants were reported in the literature for their association with PR interval (variants identified by studies in Europeans, Africans, Asians or combined ancestries)[16-23] at the start of this study. We calculated pairwise linkage disequilibrium (LD) in PLINK using the 1000 genomes (1000G) phase 3 all ancestry samples (N = 2,504)[24] for all variants within a 4Mb region centered on each previously reported variant using PLINK v1.9[25]. Multiple variants in LD ($r^2 > 0.1$) with each previously reported variant were ordered according to their positions on the chromosome, and a window was defined with the start position of the first variant in the ordered list and the end position of the last variant in the ordered list. The start and end of the window was then extended by 50kb on either side. This LD defined window or a window of ±500kb, which ever was the larger, was considered as a previously reported locus. Overlapping loci were merged and previously reported variants within the merged loci were ordered by their chromosomal position to define the start and end of the merged locus. Following this approach the 98 variants were grouped into 64 loci which we refer to as previously reported.

We followed a similar approach to define novel loci. For each lead variant within a 1Mb region outside previously reported loci we calculated pairwise LD using the 1000G phase 3 all ancestry samples (N = 2,504) or European only (N = 503)[24] within a 4Mb region. Variants in LD ($r^2 > 0.1$) with each novel variant were ordered according to their positions on the chromosome, and a window was defined with the start position of the first variant in the ordered list and the end position of the last variant in the ordered list. The start and end of the window was then extended by 50kb on either side. This LD defined window or a window of 1Mb, whichever was the larger, was considered as a novel locus. We merged overlapping loci and then ordered variants within the merged loci by their position to define the start and end of the merged locus.

### 3. Supplementary Figures

**Supplementary Figure 1** Quantile-Quantile plots of autosomal variant results from meta-analyses of PR interval.

P values from multi-ancestry (a), European (b), African (c), and Hispanic/Latino ancestry (d) meta-analyses of autosomal variants for absolute PR interval (ms) are plotted on the $-\log_{10}$ scale for all variants. The genomic inflation factors ($\lambda_{GC}$) are provided at the top left of each plot.

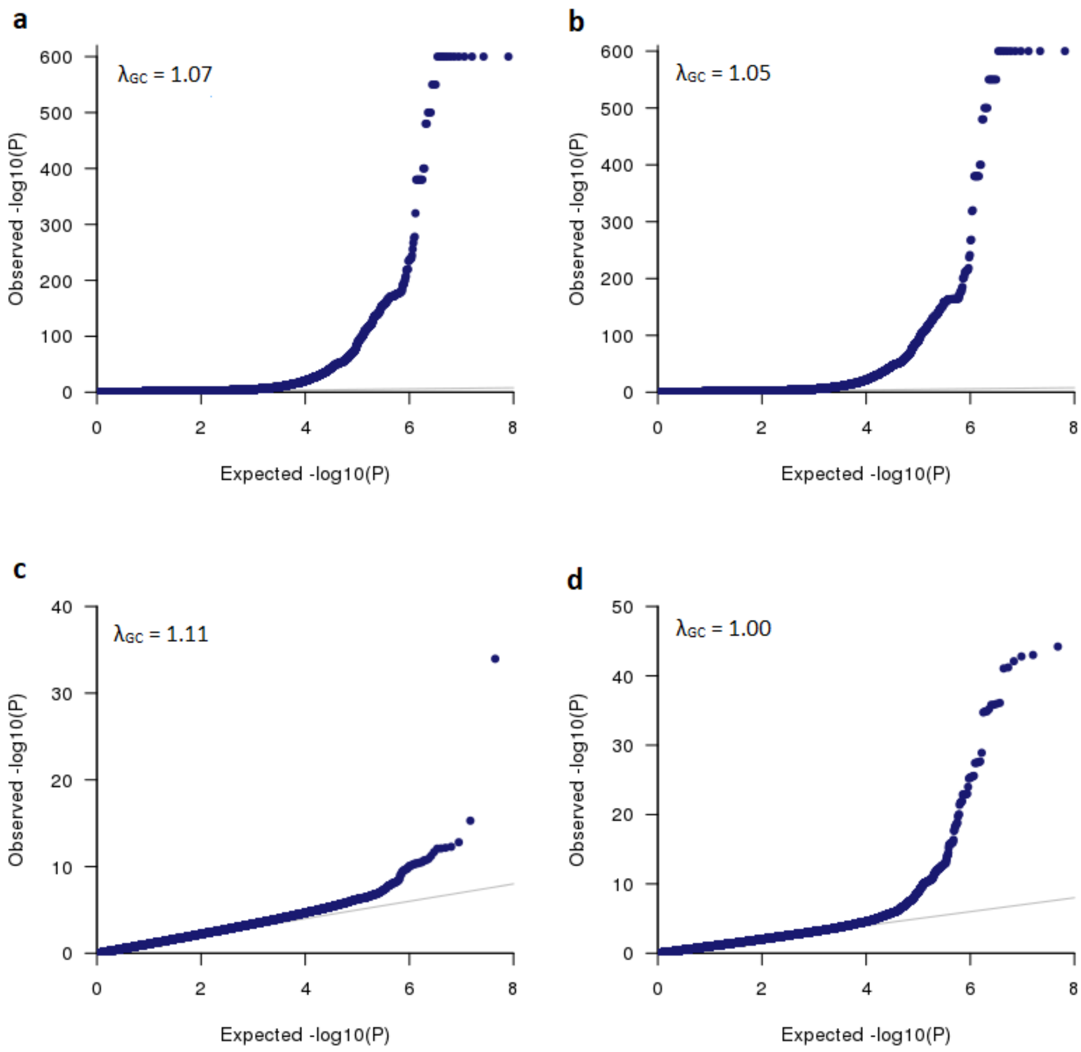

**Supplementary Figure 2** Region plots of the 149 novel loci [in a separate file].

**Supplementary Figure 3** Manhattan plot from the European (a), African (b), and Hispanic (c) meta-analyses of absolute PR interval.

P values are plotted on the $-\log_{10}$ scale for all variants present in at least 60% of the maximum sample size. Associations of genome-wide significant ($P < 5\times10^{-8}$) variants at novel and previously reported loci for PR interval are plotted in dark and light blue colors, respectively. In European meta-analysis 188 loci, of which 127 are newly identified and 61 were previously reported, reached genome-wide significance (Supplementary Tables 4, 5). Of the 127 novel loci, 119 were also genome-wide significant in the multi-ancestry meta-analysis of PR interval (Table 1, Supplementary Table 5). Eight loci exceeded the genome-wide significance threshold in African meta-analysis, of which three were not previously reported and were not observed in multi-ancestry or European meta-analyses (Supplementary Table 8). The four genome-wide significant loci that were identified by the Hispanic meta-analysis were previously reported (Supplementary Table 8).

(a)

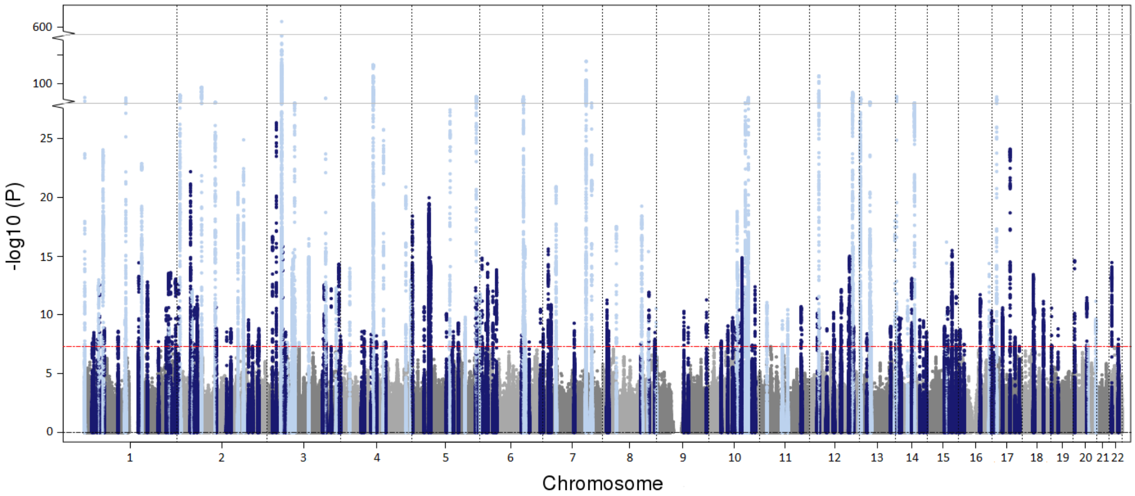


(b)

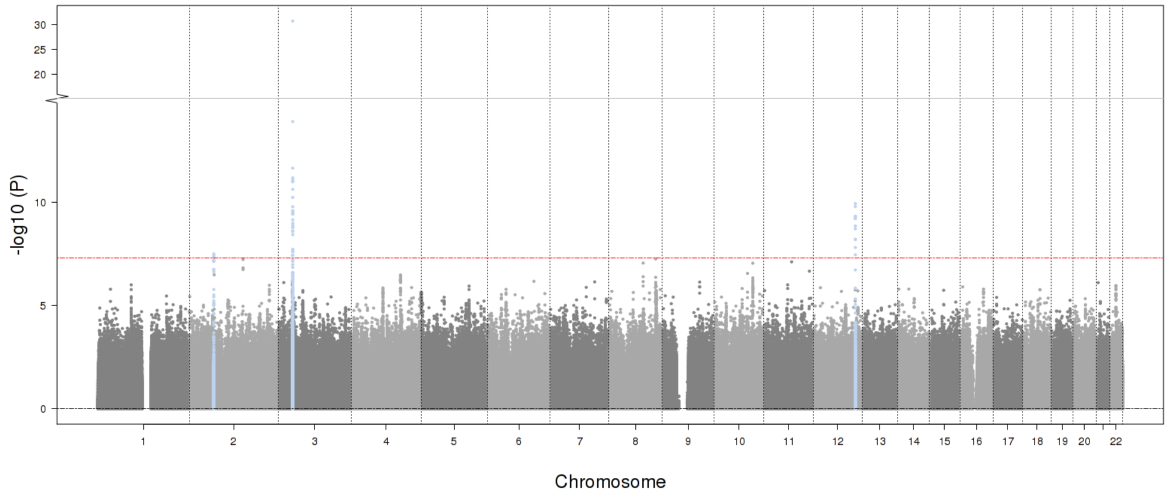

-log10 (P)
Chromosome
1 2 3 4 5 6 7 8 9 10 11 12 13 14 15 16 17 18 19 20 22


(c)

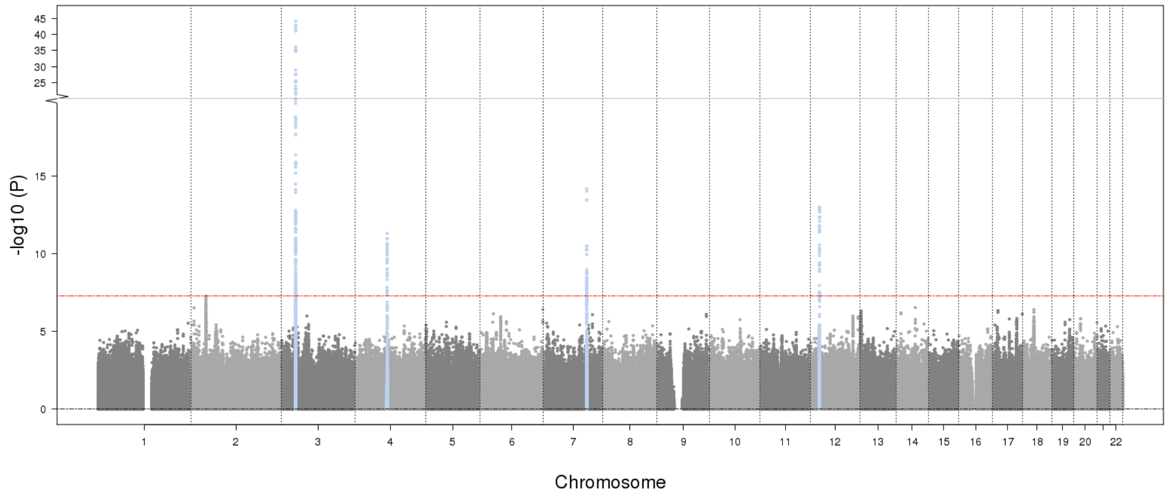

-log10 (P)
Chromosome
1 2 3 4 5 6 7 8 9 10 11 12 13 14 15 16 17 18 19 20 22

**Supplementary Figure 4** Correlation of association statistics for the lead variants at the 149 novel loci identified by the multi-ancestry and European ancestry meta-analyses of absolute PR interval.

Correlations are presented for P values (a), betas (b), and effect allele frequencies (EAF, c). Lead variants discovered from the multi-ancestry (N = 141) and European (N = 8) meta-analyses are shown in dark blue circles and light blue squares, respectively. P values are plotted on the -$\log_{10}$ scale. Red dashed lines indicate the genome-wide significance level ($P = 5 \times 10^{-8}$).

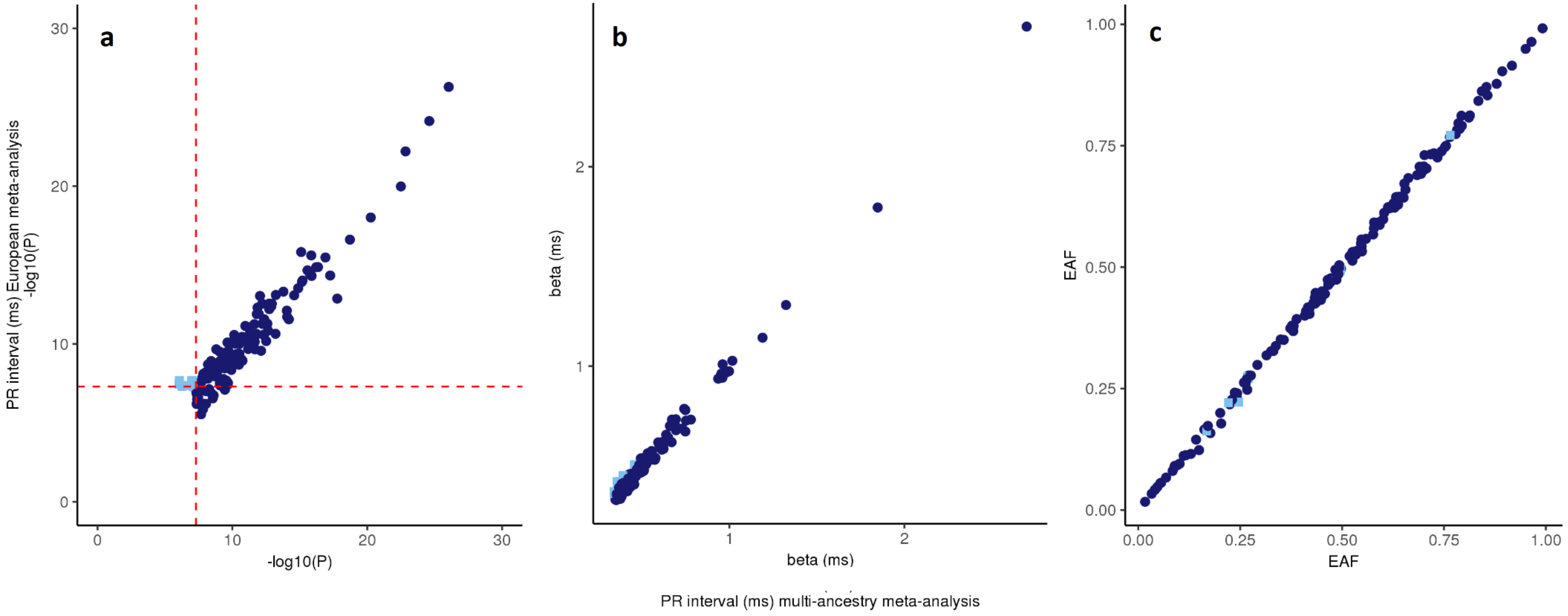

**Supplementary Figure 5** Quantile-Quantile plots of chromosome X variant results from meta-analyses of absolute PR interval.

P values from multi-ancestry (a, b), European ancestry (c, d), and African (e, f) ancestry meta-analyses of chromosome X variants for absolute PR interval (ms) are plotted on the $-\log_{10}$ scale for all variants. Chromosome X meta-analyses were performed separately for males (a, c, e) and females (b, d, f). The genomic inflation factors ($\lambda_{GC}$) are provided at the top left of each plot.

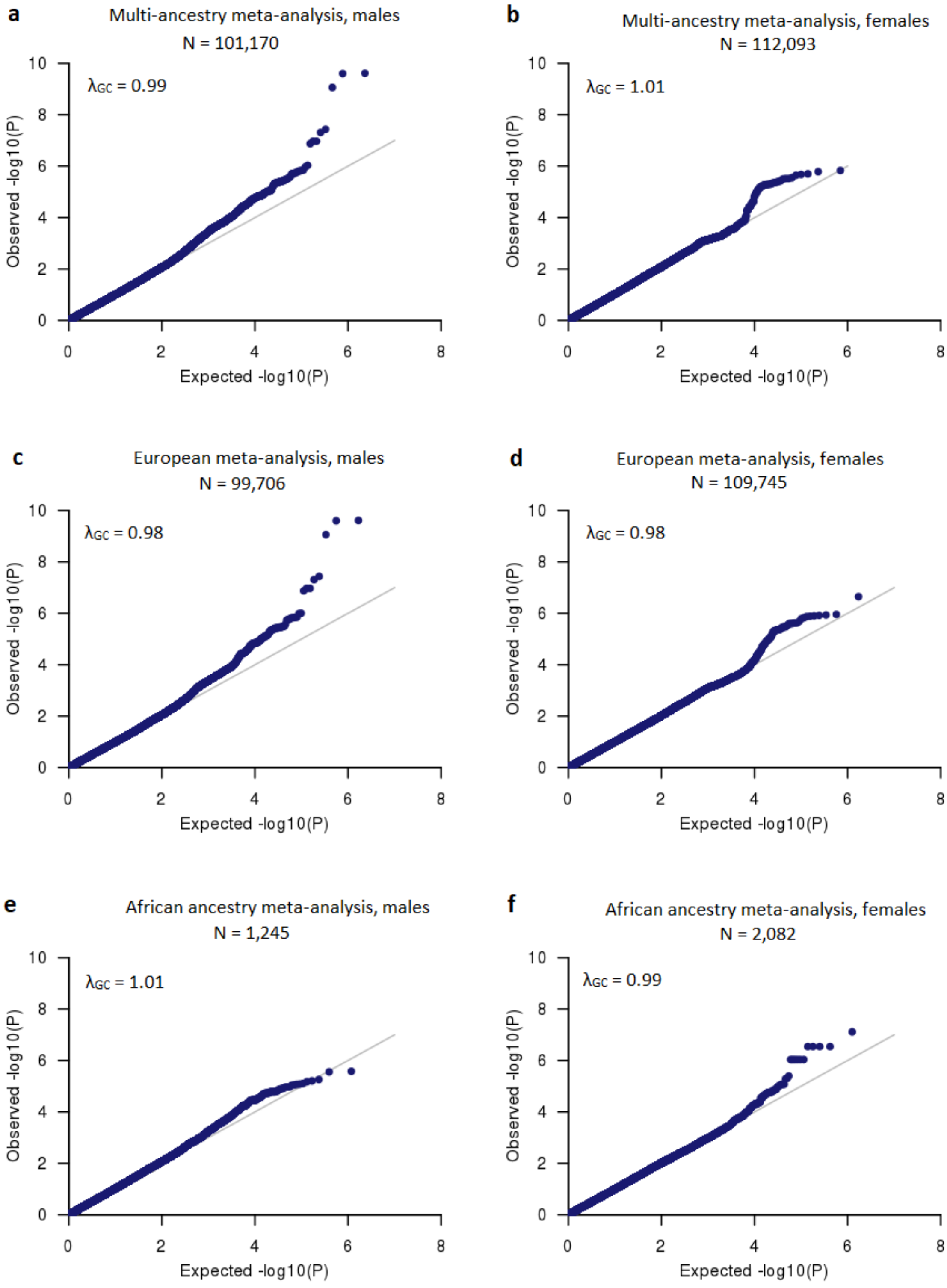
a
Multi-ancestry meta-analysis, males
N = 101,170
λGC = 0.99
b
Multi-ancestry meta-analysis, females
N = 112,093
λGC = 1.01
c
European meta-analysis, males
N = 99,706
λGC = 0.98
d
European meta-analysis, females
N = 109,745
λGC = 0.98
e
African ancestry meta-analysis, males
N = 1,245
λGC = 1.01
f
African ancestry meta-analysis, females
N = 2,082
λGC = 0.99
Observed -log10(P)
Expected -log10(P)

**Supplementary Figure 6** Correlation of P values across the meta-analyses of absolute and rank-based inverse normal transformed residuals of PR interval.

Correlations across of P values across multi-ancestry and European ancestry meta-analyses of absolute PR interval and rank-based inverse normal transformed PR interval are presented. P values are plotted for variants with $P < 10^{-6}$ in any of the four meta-analyses outside previous reported loci (N = 16,267) that were available in at least 60% of the maximum sample size of each meta-analysis on the $-\log_{10}$ scale. The blue lines show the corresponding regression lines. Red dashed lines indicate the genome-wide significance level ($P = 5 \times 10^{-8}$). Correlation coefficients ($\rho$) between the two P values are provided on the top left of each plot.

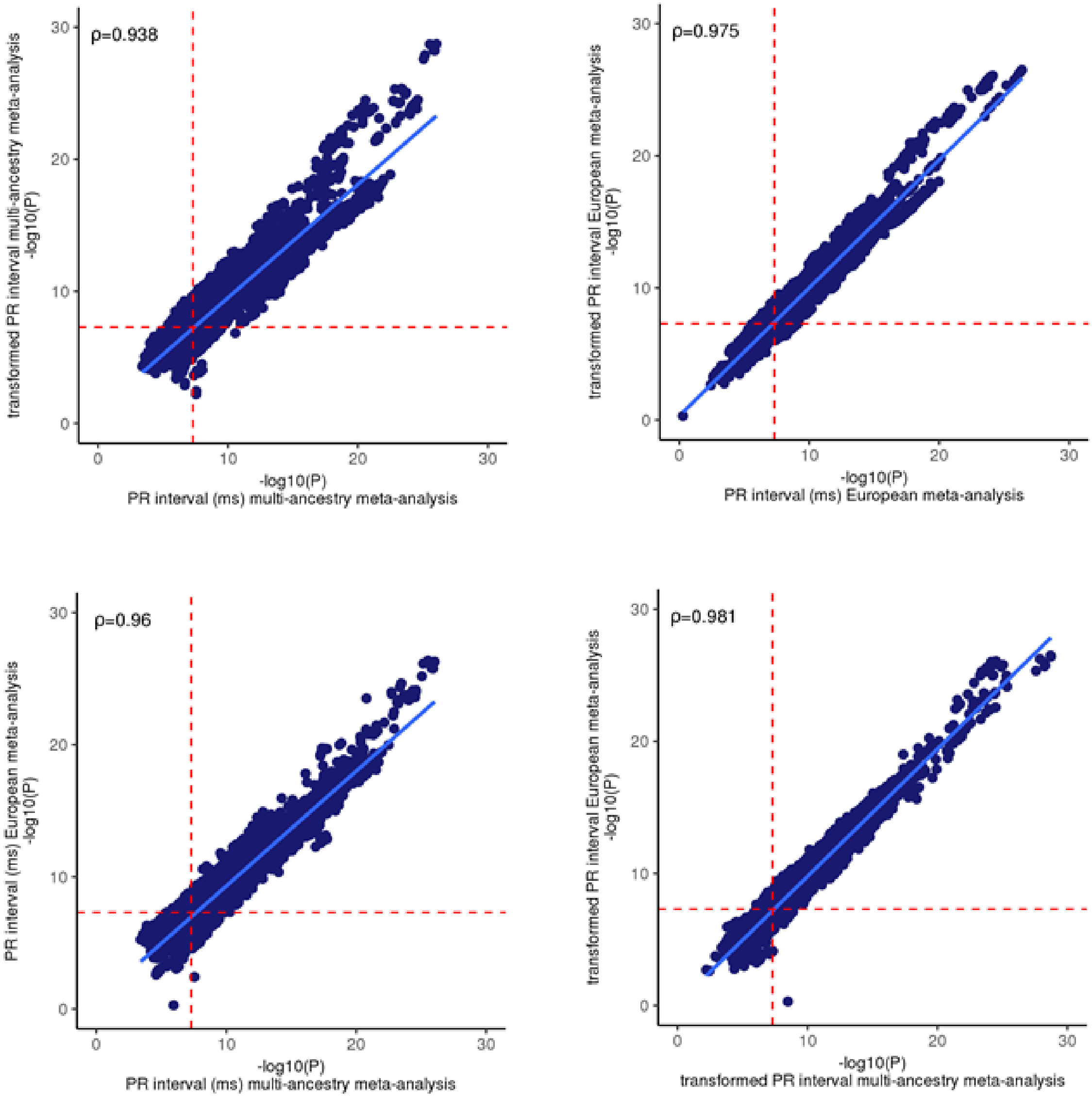
ρ=0.938
transformed PR interval multi-ancestry meta-analysis -log10(P)
-log10(P) PR interval (ms) multi-ancestry meta-analysis
ρ=0.975
transformed PR interval European meta-analysis -log10(P)
-log10(P) PR interval (ms) European meta-analysis
ρ=0.96
PR interval (ms) European meta-analysis -log10(P)
-log10(P) PR interval (ms) multi-ancestry meta-analysis
ρ=0.981
transformed PR interval European meta-analysis -log10(P)
-log10(P) transformed PR interval multi-ancestry meta-analysis

**Supplementary Figure 7** Genome-wide significant loci ($P = 5 \times 10^{-8}$) across meta-analyses of absolute and rank-based inverse normal transformed PR interval.

The Venn diagram shows overlap of genome-wide significant loci ($P = 5 \times 10^{-8}$) across multi-ancestry and European ancestry meta-analyses of absolute (darker blue circles; N = 293,051 and N = 271,570 respectively) and rank-based inverse normal transformed PR interval (light blue circles; N = 282,128 and N = 271,570 respectively). A total of 130 out of the 149 novel loci discovered by the meta-analyses of absolute PR interval reached also genome-wide significance in the meta-analyses of rank-based inverse normal transformed PR interval.

* One locus reached genome-wide significance in multi-ancestry meta-analysis of absolute PR interval and in European meta-analysis of rank-based inverse normal transformed PR interval.

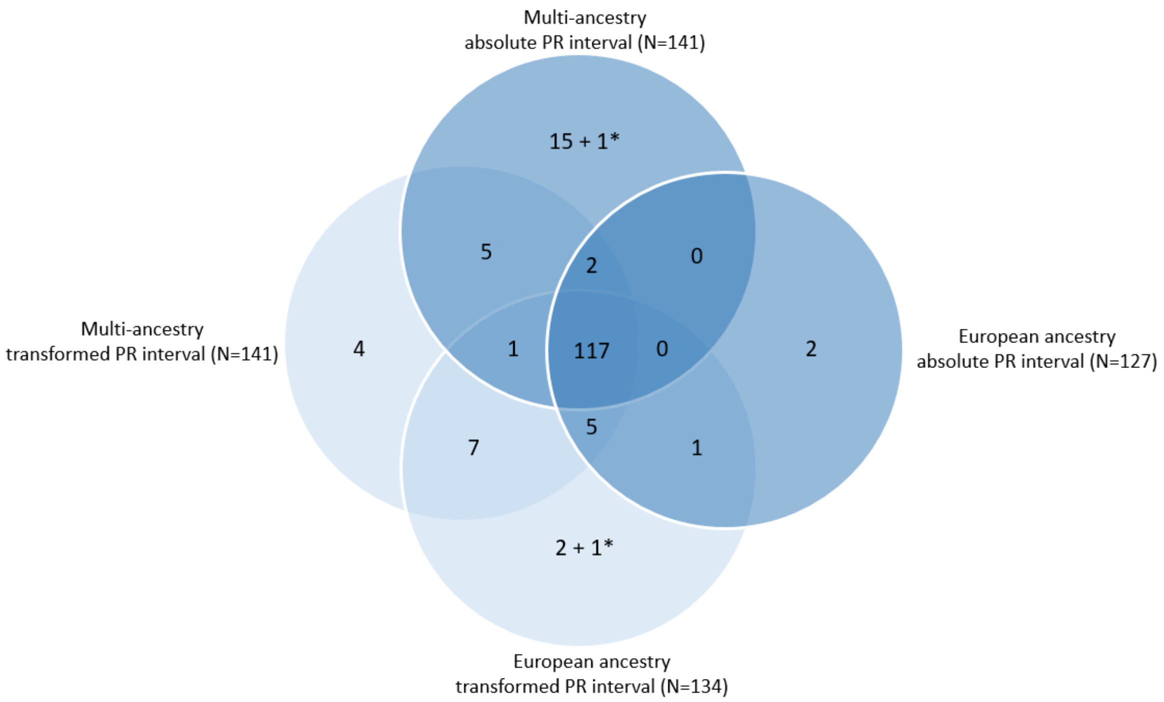

**Supplementary Figure 8** Volcano plot of predicted gene expression analysis in cardiac tissues.

The plots show the results from predicted gene expression analysis in left ventricle (a) and right atrial appendage (b) tissues from GTEx. Analysis was performed with S-PrediXcan using the European meta-analysis summary level results. The x-axis shows the effect size for associations of predicted gene expression and PR interval duration for each gene. The y-axis shows the $-\log_{10}(P)$ for the associations per gene. Each plotted point represents the association results of a single gene. The highlighted genes are significant after Bonferroni correction for all tested genes at the two tissues with a $P < 4.4 \times 10^{-6}$ (=0.05/(5,977+5,366)). Genes with positive effect (blue) showed an association of increased predicted gene expression with PR interval duration. Genes with negative effect (yellow) showed an association of decreased predicted gene expression with PR interval duration.

**a**

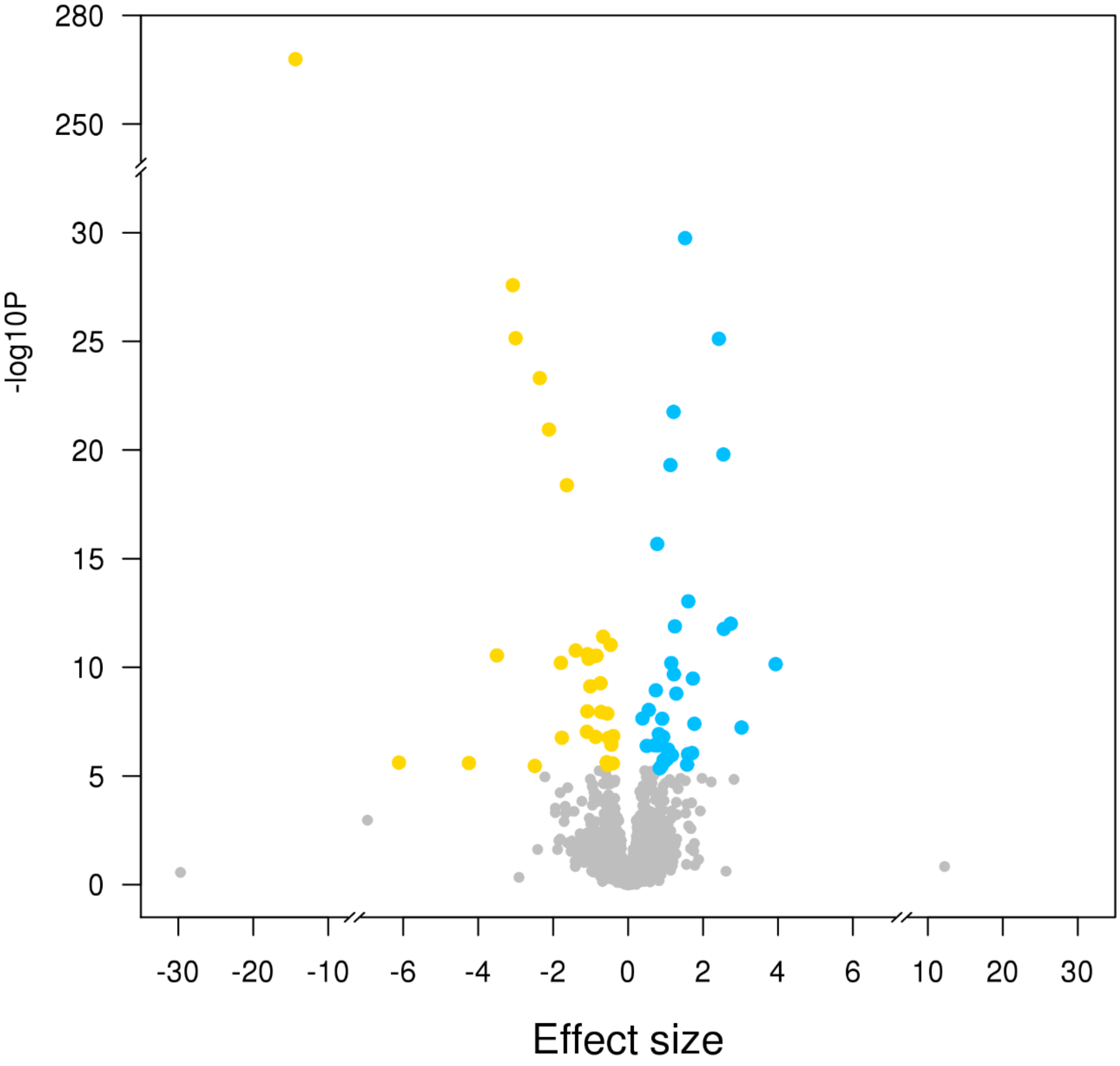
-log10P
280
250
30
25
20
15
10
5
0
-30
-20
-10
-6
-4
-2
0
2
4
6
10
20
30
Effect size

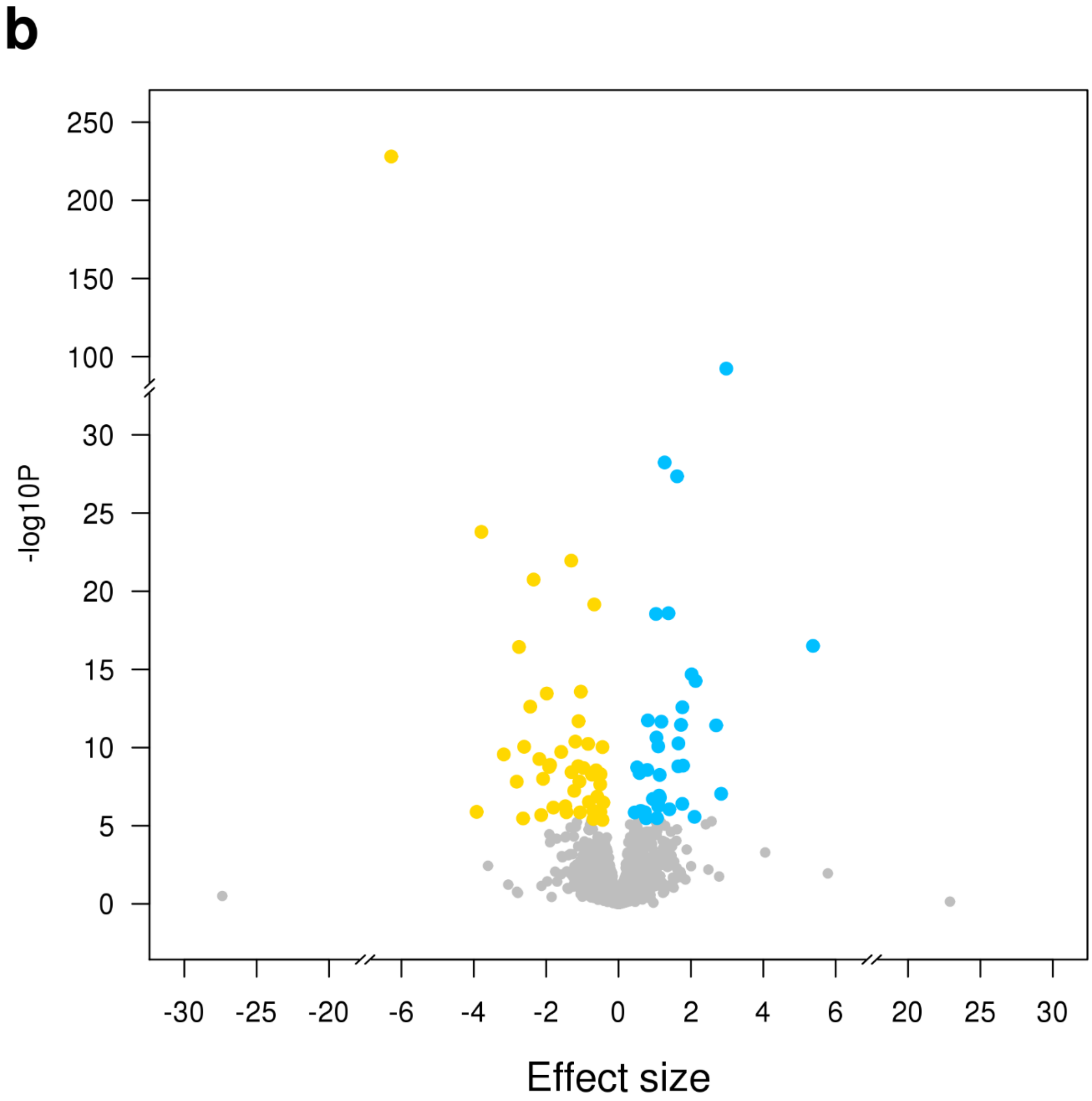
b
250
200
150
100
30
25
20
15
10
5
0
-log10P
-30
-25
-20
-6
-4
-2
0
2
4
6
20
25
30
Effect size

**Supplementary Figure 9** Enrichment of PR interval variants in DNAse I Hypersensitive sites.

Radial plot shows odds ratio (OR) values at two GWAS P-value thresholds ($T<10^{-5}$ and $T<10^{-8}$, shown by inner colors and bottom legend) for all ENCODE[28] and Roadmap Epigenomics[29] DHS cell lines, sorted by tissue on the outer circle. Small dots on the outer side of the plot show significant enrichment (if present) at $T<10^{-5}$ (outermost) to $T<10^{-8}$ (innermost) after multiple-testing correction for the number of effective annotations and are colored with respect to the tissue cell type tested (font size of tissue labels reflects the number of cell types from that tissue).

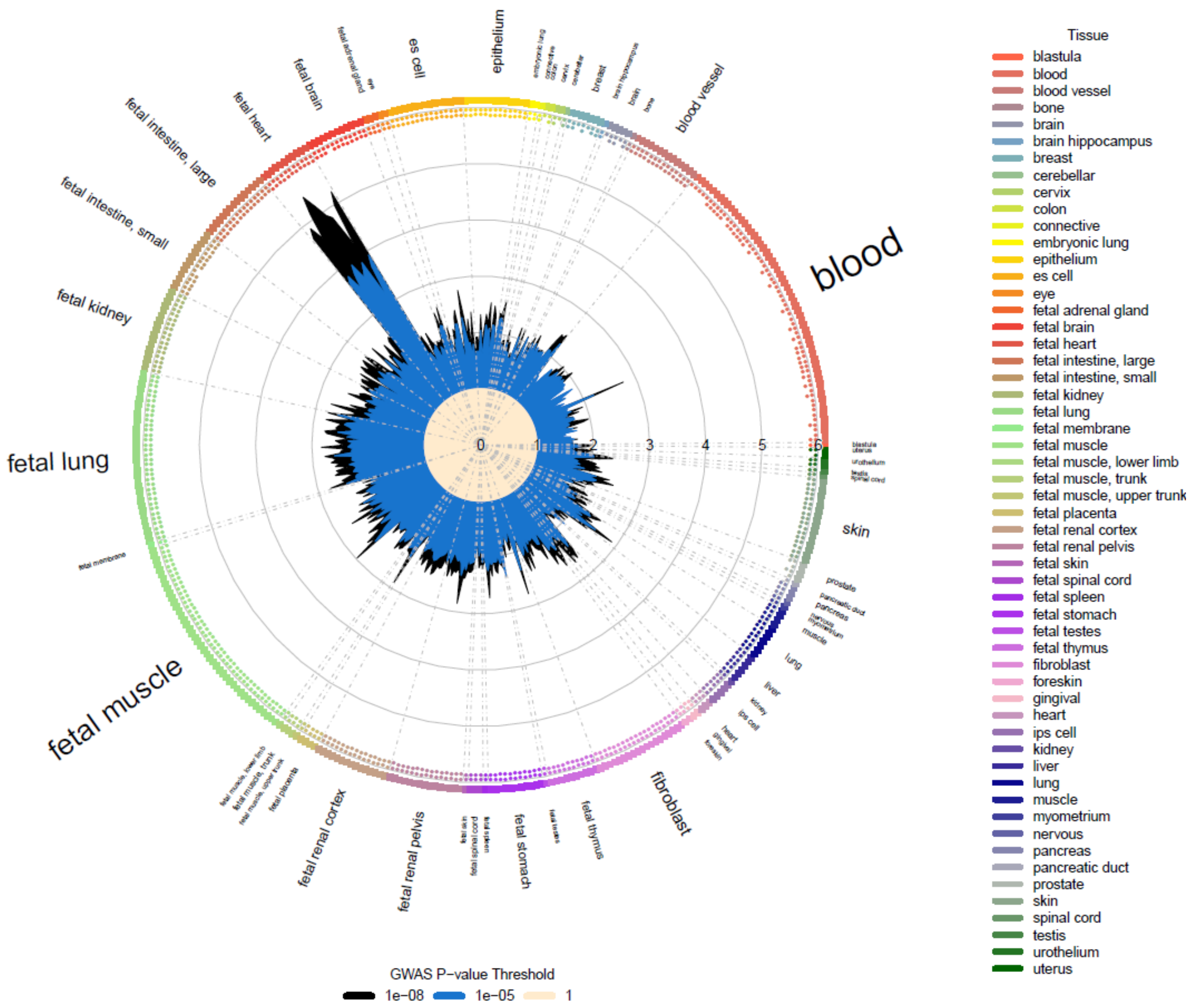

**Supplementary Figure 10** Association of novel PR interval loci with other GWAS traits.

The chord plot shows results for genome-wide significant ($P < 5\times10^{-8}$) associations with other GWAS traits which were extracted from PhenoScanner v2[26] database for the lead and conditionally independent variants at the 149 novel loci (Table 1, Supplementary Table 11) as well as their proxies ($r^2>0.8$). Traits were grouped into broader categories, and results are presented only for traits associated with three or more PR interval loci. Detailed PhenoScanner GWAS results are presented in Supplementary Table 17. For atrial fibrillation, we curated a list of all overlapping loci ($r^2>0.7$) with PR interval including the recent findings from two GWASs[2,27] not included in PhenoScanner v2 database.

ECG, electrocardiogram; CAD: coronary artery disease; CHD: coronary heart disease; BMI, body mass index; WC, waist circumference; HC: hip circumference.

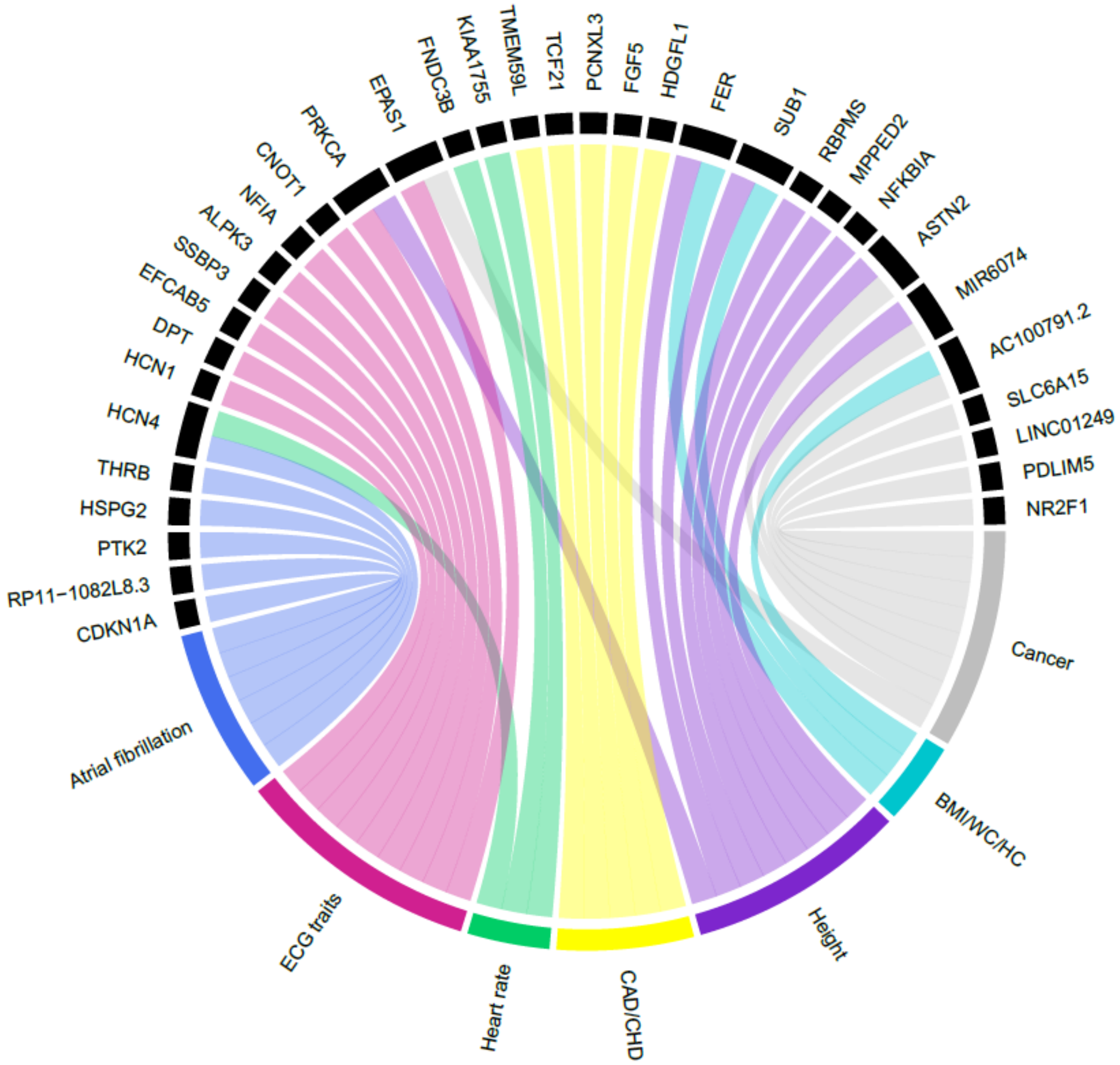

CDKN1A
RP11−1082L8.3
PTK2
HSPG2
THRB
HCN4
HCN1
DPT
EFCAB5
SSBP3
ALPK3
NFIA
CNOT1
PRKCA
EPAS1
FNDC3B
KIAA1755
TMEM59L
TCF21
PCNXL3
FGF5
HDGFL1
FER
SUB1
RBPMS
MPPED2
NFKBIA
ASTN2
MIR6074
AC100791.2
SLC6A15
LINC01249
PDLIM5
NR2F1
Cancer
BMI/WC/HC
Height
CAD/CHD
Heart rate
ECG traits
Atrial fibrillation


644

645

**Supplementary Figure 11** Bubble plot of phenome-wide association analysis of European-ancestry PR interval polygenic risk score.

Polygenic risk score derived from the European-ancestry meta-analysis results. Orange circles indicate that higher polygenic risk of prolonged PR interval is associated with an increased risk of the condition, whereas blue circles indicate that higher polygenic risk of prolonged PR interval is associated with a lower risk of the condition. The darkness of the color reflects the odds ratio (OR) changes per 1 standard deviation increment of the polygenic risk score. Given correlation between traits, we did not establish a pre-specified significance threshold for the analysis and report nominal associations ($P < 0.05$).

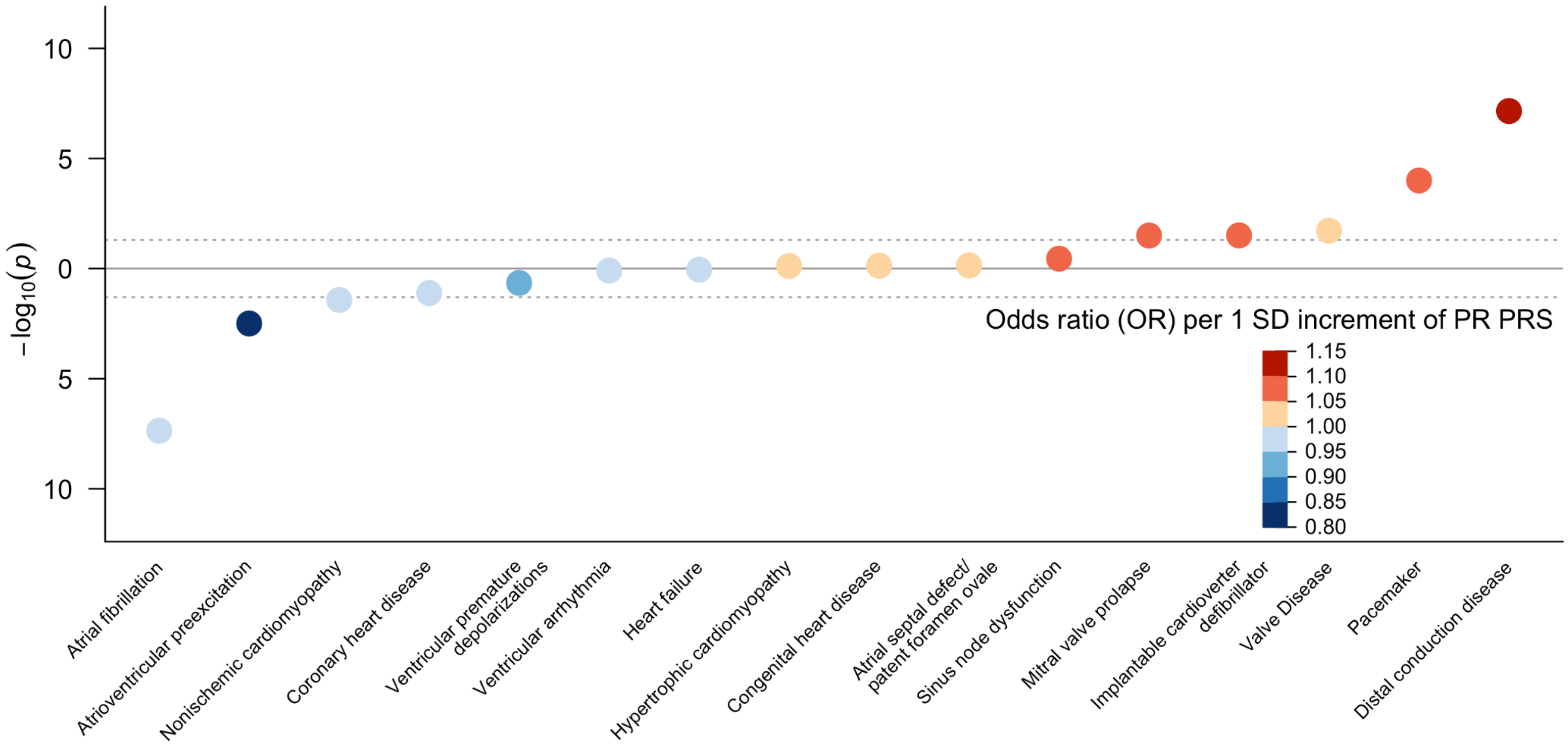
−log10(p)
10
5
0
5
10
Odds ratio (OR) per 1 SD increment of PR PRS
1.15
1.10
1.05
1.00
0.95
0.90
0.85
0.80
Atrial fibrillation
Atrioventricular preexcitation
Nonischemic cardiomyopathy
Coronary heart disease
Ventricular premature depolarizations
Ventricular arrhythmia
Heart failure
Hypertrophic cardiomyopathy
Congenital heart disease
Atrial septal defect/ patent foramen ovale
Sinus node dysfunction
Mitral valve prolapse
Implantable cardioverter defibrillator
Valve Disease
Pacemaker
Distal conduction disease

### 4. Acknowledgements

**AMISH**

We gratefully thank our Amish community and research volunteers for their long-standing partnership in research, and acknowledge the dedication of our Amish liaisons, field workers and the Amish Research Clinic staff, without which these studies would not have been possible.

**ARIC**

The authors thank the staff and participants of the ARIC study for their important contributions.

**BioME**

The Mount Sinai BioMe Biobank is supported by The Andrea and Charles Bronfman Philanthropies.

**BRIGHT**

The BRIGHT study is extremely grateful to all the patients who participated in the study and the BRIGHT nursing team and staff at the Genome Centre for genotyping. This work forms part of the research program of the National Institutes of Health Research (NIHR Cardiovascular Biomedical Research) Cardiovascular Biomedical Centre at Barts and The London, QMUL.

**CHRIS**

Full acknowledgements for the CHRIS study are reported here: http://translational-medicine.biomedcentral.com/articles/10.1186/s12967-015-0704-9#Declarations. The CHRIS study was funded by the Department of Innovation, Research, and University of the Autonomous Province of Bolzano-South Tyrol.

**CROATIA-Korcula**

We would like to acknowledge the contributions of the recruitment team in Korcula, the administrative teams in Croatia and Edinburgh and the people of Korcula. The SNP genotyping for the KORCULA cohort was performed in Helmholtz Zentrum München, Neuherberg, Germany and the Clinical Research Facility Genetics Core, University of Edinburgh at the Western General Hospital, Edinburgh, UK.

**CROATIA-Split**

We would like to acknowledge the contributions of the recruitment team from the Croatian Centre for Global Health, University of Split, the administrative teams in Croatia and Edinburgh and the people of Split. The SNP genotyping for the CROATIA_Split cohort was performed by AROS Applied Biotechnology, Aarhus, Denmark and the Clinical Research Facility Genetics Core, University of Edinburgh at the Western General Hospital, Edinburgh, UK.

**deCODE**

The deCODE researchers gratefully thank the study participants, the staff at deCODE and our recruitment center, and all our collaborators for invaluable contributions to the understanding of the genetics of health and disease.

**ERF**

We are grateful to all study participants and their relatives, general practitioners and neurologists for their contributions to the ERF study and to P Veraart for her help in genealogy, J Vergeer for the supervision of the laboratory work and P Snijders for his help in data collection.

**FHS**

We thank the FHS participants and staff for their contributions to this work.

**FINCAVAS**

The authors thank the staff of the Department of Clinical Physiology for collecting the exercise test data.

**GAPP**

We thank the GAPP staff and all GAPP study participants for their important contributions

**GRAPHIC**

This work falls under the portfolio of research supported by the NIHR Leicester Biomedical Research Centre.

**GS:SFHS**

We are grateful to all the families who took part, the general practitioners and the Scottish School of Primary Care for their help in recruiting them, and the whole Generation Scotland team, which includes interviewers, computer and laboratory technicians, clerical workers, research scientists, volunteers, managers, receptionists, healthcare assistants and nurses.

**HCHS/SOL**

We thank the participants and staff of the HCHS/SOL study for their contributions to this study.

**Health 2000**

The Health 2000 Survey is extremely grateful to all the participants of the study.

**INGI-CARL/ INGI-FVG**

We are very grateful to the municipal administrators for their collaboration on the project and for logistic support. We would like to thank all participants to this study.

**JHS**

We thank the Jackson Heart Study (JHS) participants and staff for their contributions to this work.

**KORA F3/ KORA S4**

The KORA study was initiated and financed by the Helmholtz Zentrum München – German Research Center for Environmental Health, which is funded by the German Federal Ministry of Education and Research (BMBF) and by the State of Bavaria. Furthermore, KORA research was supported within the Munich Center of Health Sciences (MC-Health), Ludwig-Maximilians-Universität, as part of LMUinnovativ.

**Lifelines**

We thank U. Bultmann (1), J.M. Geleijnse (2), P. van der Harst (3), S. Mulder (4), J.G.M. Rosmalen (5), E.F.C. van Rossum (6), H.A. Smit (7), M. Swertz (8), E.A.L.M. Verhagen (9)

(1) Department of Social Medicine, University Medical Center Groningen, University of Groningen, The Netherlands, (2) Department of Human Nutrition, Wageningen University, The Netherlands, (3) Department of Cardiology, University Medical Center Groningen, University of Groningen, The Netherlands, (4) Lifelines Cohort Study, The Netherlands, (5) Interdisciplinary Center of Psychopathology of Emotion Regulation (ICPE), Department of Psychiatry, University Medical Center Groningen, University of Groningen, The Netherlands, (6) Department of Endocrinology, Erasmus Medical Center, Rotterdam, The Netherlands, (7) Department of Public Health, University Medical Center Utrecht, The Netherlands, (8)

Department of Genetics, University Medical Center Groningen, University of Groningen, The Netherlands, (9) Department of Public and Occupational Health, VU Medical Center, Amsterdam, The Netherlands

**MICROS**

We owe a debt of gratitude to all participants, all primary care practitioners, and the personnel of the Hospital of Silandro (Department of Laboratory Medicine) for their participation and collaboration in the research project. The study was supported by the Ministry of Health and Department of Educational Assistance, University and Research of the Autonomous Province of Bolzano, the South Tyrolean Sparkasse Foundation, and the European Union framework program 6 EUROSPAN project (contract no. LSHG-CT-2006-018947).

**NEO**

The authors of the NEO study thank all individuals who participated in the Netherlands Epidemiology in Obesity study, all participating general practitioners for inviting eligible participants and all research nurses for collection of the data. We thank the NEO study group, Pat van Beelen, Petra Noordijk and Ingeborg de Jonge for the coordination, lab and data management of the NEO study. We also thank Arie Maan for the analyses of the electrocardiograms.

**ORCADES**

DNA extractions were performed at the Wellcome Trust Clinical Research Facility in Edinburgh. We would like to acknowledge the invaluable contributions of the research nurses in Orkney, the administrative team in Edinburgh and the people of Orkney.

**PIVUS**

The investigators express their deepest gratitude to the study participants.

**PROSPER**

The PROSPER study was supported by an investigator initiated grant obtained from Bristol-Myers Squibb. Prof. Dr. J. W. Jukema is an Established Clinical Investigator of the Netherlands Heart Foundation (grant 2001 D 032). Support for genotyping was provided by the seventh framework program of the European commission (grant 223004) and by the Netherlands Genomics Initiative (Netherlands Consortium for Healthy Aging grant 050-060-810).

**RS**

We thank Pascal Arp, Mila Jhamai, Dr Michael Moorhouse, Marijn Verkerk, and Sander Bervoets for their help in creating the GWAS database. The authors are very grateful to the participants and staff from the Rotterdam Study, the participating general practitioners and the pharmacists.

**SardiNIA**

We thank the many volunteers who generously participated in this study, the Mayors and citizens of the Sardinian towns involved, the head of the Public Health Unit ASL4, and the province of Ogliastra for their volunteerism and cooperation. In addition, we are grateful to the Mayor and the administration in Lanusei for providing and furnishing the clinic site. We are grateful to the physicians Angelo Scuteri, Marco Orrù, Maria Grazia Pilia, Liana Ferreli, Francesco Loi, nurses Paola Loi, Monica Lai and Anna Cau who carried out participant physical exams; the recruitment personnel Susanna Murino; and Mariano Dei, Sandra Lai, Andrea Maschio, Fabio Busonero for genotyping. This research was supported by the Intramural Research Program of the NIH, National Institute on Aging with contracts N01-AG-1-2109 and HHSN271201100005C, and by grant FaReBio2011 “Farmaci e Reti Biotecnologiche di Qualità”.

**UK Biobank**

This research has been conducted using the UK Biobank Resource (application 8256 - Understanding genetic influences in the response of the cardiac electrical system to exercise). This work forms part of the research program of the National Institutes of Health Research (NIHR Cardiovascular Biomedical Research) Cardiovascular Biomedical Centre at Barts and The London, QMUL. PD Lambiase acknowledges support from the UCLH Biomedicine NIHR.

Polygenic risk score analyses were also conducted using the UK Biobank Resource (application 17488 - Understanding the Genetic Basis of Cardiac Arrhythmias).

**ULSAM**

The investigators express their deepest gratitude to the study participants.

**WHI**

The WHI program is supported by contracts from the National Heart, Lung and Blood Institute, NIH. The authors thank the WHI investigators and staff for their dedication, and the study participants for making the program possible. All contributors to WHI science are listed at https://www.whi.org/researchers/Documents%20%20Write%20a%20Paper/WHI%20Investigator%20Long%20List.pdf.https://www.whi.org/researchers/Documents%20%20Write%20a%20Paper/WHI%20Investigator%20Short%20List.pdf.

**YFS**

We thank the teams that collected data at all measurement time points; the persons who participated as both children and adults in these longitudinal studies; and biostatisticians Irina Lisinen, Johanna Ikonen, Noora Kartiosuo, Ville Aalto, and Jarno Kankaanranta for data management and statistical advice.

**Additional acknowledgements**

This research utilized Queen Mary's Apocrita HPC facility, supported by QMUL Research-IT. http://doi.org/10.5281/zenodo.438045

Figure 5 was designed using BioRender software (biorender.io).

### 5. Study funding

**AMISH**

The Amish studies are supported by grants and contracts from the NIH, including R01 AG18728, R01 HL088119, U01 GM074518, U01 HL072515, U01 HL84756, U01 HL137181, R01 DK54261,the University of Maryland General Clinical Research Center, grant M01 RR 16500, and the Mid-Atlantic Nutrition Obesity Research Center grant P30 DK72488, the Baltimore Diabetes Research and Training Center grant P60DK79637.

**ARIC**

The Atherosclerosis Risk in Communities Study is carried out as a collaborative study supported by National Heart, Lung, and Blood Institute contracts (HHSN268201100005C, HHSN268201100006C, HHSN268201100007C, HHSN268201100008C, HHSN268201100009C, HHSN268201100010C, HHSN268201100011C, and HHSN268201100012C), R01HL087641, R01HL59367 and R01HL086694; National Human Genome Research Institute contract U01HG004402; and National Institutes of Health contract HHSN268200625226C. Infrastructure was partly supported by Grant Number UL1RR025005, a component of the National Institutes of Health and NIH Roadmap for Medical Research.

**BAMBUÍ**

Ministry of Science and Technology (Financiadora de Estudos e Projetos – FINEP and Conselho Nacional de Desenvolvimento Científico e Tecnologico – CNPq), and Fundação de Amparo a Pesquisa do Estado de Minas Gerais (FAPEMIG).

**BRIGHT**

This work was supported by the Medical Research Council of Great Britain (grant number G9521010D) and the British Heart Foundation (grant number PG/02/128).

**Broad-AF**

Dr. Lubitz is supported by NIH grant 1R01HL139731 and American Heart Association 18SFRN34250007. Dr. Fatkin is supported by the Victor Chang Cardiac Research Institute, National Health and Medical Research Council of Australia, Estate of the Late RT Hall, St Vincent's Clinic Foundation, and the Simon Lee Foundation. Dr. Olesen is supported by the Hallas-Møller Emerging Investigator Novo Nordisk (NNF17OC0031204). Dr. Smith was supported by grants from the Swedish Heart-Lung Foundation (2016-0134 and 2016-0315), the Swedish Research Council (2017-02554), the European Research Council (ERC-STG-2015-679242), the Crafoord Foundation, Skåne University Hospital, the Scania county, governmental funding of clinical research within the Swedish National Health Service, a generous donation from the Knut and Alice Wallenberg foundation to the Wallenberg Center for Molecular Medicine in Lund, and funding from the Swedish Research Council (Linnaeus grant Dnr 349-2006-237, Strategic Research Area Exodiab Dnr 2009-1039) and Swedish Foundation for Strategic Research (Dnr IRC15-0067) to the Lund University Diabetes Center. Dr. Weng is supported by the American Heart Association Postdoctoral Fellowship Award 17POST33660226. This work was also supported by an American Heart Association Strategically Focused Research Networks (SFRN) postdoctoral fellowship to Drs. Weng and Hall (18SFRN34110082).

This work was supported by grants from the National Institutes of Health to Dr. Ellinor (1RO1HL092577, R01HL128914, K24HL105780). Dr. Ellinor is also supported by the American Heart Association (18SFRN34110082) and by the Fondation Leducq (14CVD01). This research was also supported by the Carol and Roch Hillenbrand and the George L. Nardi, MD, funds at Massachusetts General Hospital. The GGAF is supported by funding to the 5 sources that form GGAF. The AF RISK study is supported by the Netherlands Heart Foundation (grant NHS2010B233), and the Center for Translational Molecular Medicine. Both the Young-AF and Biomarker-AF studies are supported by funding from the University Medical Center Groningen. The GIPS-III trial was supported by grant 95103007 from ZonMw, the Netherlands Organization for Health Research and Development. The PREVEND study is supported by the Dutch Kidney Foundation (grant E0.13) and the Netherlands Heart Foundation (grant NHS2010B280). Dr. Rienstra acknowledges support from the Netherlands Cardiovascular Research Initiative: an initiative supported by the Netherlands Heart Foundation, CVON 2014–9: "Reappraisal of Atrial Fibrillation: interaction between hyperCoagulability, Electrical remodelling, and Vascular destabilization in the progression of AF (RACE V). The Intermountain was supported by funding from the Dell Loy Hansen Heart Foundation. The re-examination of the MPP study was supported by The Swedish Heart-Lung Foundation, Merck, Sharp & Dohme, Hulda and E Conrad Mossfelts Foundation and Ernhold Lundströms Foundation. J. Gustav Smith was supported by the European Research Council, Swedish Heart-Lung Foundation, the Wallenberg Center for Molecular Medicine at Lund University, the Swedish Research Council, the Crafoord Foundation, governmental funding of clinical research within the Swedish National Health Service, Skåne University Hospital in Lund, and the Scania county. Vanderbilt University Medical Center's BioVU projects are supported by numerous sources: institutional funding, private agencies, and federal grants. These include the NIH funded Shared Instrumentation Grant S10RR025141; CTSA grants

UL1TR002243, UL1TR000445, and UL1RR024975. Genomic data are also supported by investigator-led projects that include U01HG004798, R01NS032830, RC2GM092618, P50GM115305, U01HG006378, U19HL065962, R01HD074711; and additional funding sources listed at https://victr.vanderbilt.edu/pub/biovu/?sid=229. The Danish AF Study support from the John and Birthe Meyer Foundation, The Research Foundation at Rigshospitalet, Villadsen Family Foundation, The Arvid Nilsson Foundation, and The Hallas-Møller Emerging Investigator Novo Nordisk (NNF17OC0031204).

**CHS**

Cardiovascular Health Study: This CHS research was supported by NHLBI contracts HHSN268201200036C, HHSN268200800007C, HHSN268201800001C, N01HC55222, N01HC85079, N01HC85080, N01HC85081, N01HC85082, N01HC85083, N01HC85086, and HHSN268200960009C; and NHLBI grants U01HL080295, R01HL087652, R01HL105756, R01HL103612, R01HL120393, R01HL085251, and U01HL130114 with additional contribution from the National Institute of Neurological Disorders and Stroke (NINDS). Additional support was provided through R01AG023629 from the National Institute on Aging (NIA). A full list of principal CHS investigators and institutions can be found at CHS-NHLBI.org.

The provision of genotyping data was supported in part by the National Center for Advancing Translational Sciences, CTSI grant UL1TR001881, and the National Institute of Diabetes and Digestive and Kidney Disease Diabetes Research Center (DRC) grant DK063491 to the Southern California Diabetes Endocrinology Research Center.

The content is solely the responsibility of the authors and does not necessarily represent the official views of the National Institutes of Health.

**CROATIA-Korcula/CROATIA-Split**

Medical Research Council UK and the Ministry of Science, Education and Sport in the Republic of Croatia (number 108-1080315-0302).

**deCODE**

All authors affiliated with deCODE genetics/Amgen, Inc. are employed by the company

**ERF**

The ERF study as a part of EUROSPAN (European Special Populations Research Network) was supported by European Commission FP6 STRP grant number 018947 (LSHG-CT-2006-01947) and also received funding from the European Community's Seventh Framework Programme (FP7/2007-2013)/grant agreement HEALTH-F4-2007-201413 by the European Commission under the programme "Quality of Life and Management of the Living Resources" of 5th Framework Programme (no. QLG2-CT-2002-01254). The ERF study was further supported by ENGAGE consortium and CMSB. High-throughput analysis of the ERF data was supported by joint grant from Netherlands Organisation for Scientific Research and the Russian Foundation for Basic Research (NWO-RFBR 047.017.043).

**FHS**

The Framingham Heart Study is funded by National Institutes of Health contract N01-HC-25195; HHSN268201500001I. The current project was also supported by 1R01HL128914; 2R01 HL092577; 1P50HL120163 and 18SFRN34110082.

**FINCAVAS**

The Finnish Cardiovascular Study (FINCAVAS) has been financially supported by the Competitive Research Funding of the Tampere University Hospital (Grant 9M048 and 9N035), the Finnish Cultural Foundation, the Finnish Foundation for Cardiovascular Research, the Emil Aaltonen Foundation, Finland, the Tampere Tuberculosis Foundation, and EU Horizon 2020 (grant 755320 for TAXINOMISIS).

**GAPP**

The GAPP study was supported by the Liechtenstein Government, the Swiss Heart Foundation, the Swiss Society of Hypertension, the University of Basel, the University Hospital Basel, the Hanela Foundation, the Mach-Gaensslen Foundation, Schiller AG, and Novartis.

**GRAPHIC**

The GRAPHIC Study was funded by the British Heart Foundation (BHF/RG/2000004). CPN and NJS are supported by the British Heart Foundation and NJS is a NIHR Senior Investigator

**GS:SFHS**

Generation Scotland received core support from the Chief Scientist Office of the Scottish Government Health Directorates [CZD/16/6] and the Scottish Funding Council [HR03006]. Genotyping of the GS:SFHS samples was carried out by the Genetics Core Laboratory at the Wellcome Trust Clinical Research Facility, Edinburgh, Scotland and was funded by the Medical Research Council UK and the Wellcome Trust (Wellcome Trust Strategic Award "STratifying Resilience and Depression Longitudinally" (STRADL) Reference 104036/Z/14/Z).

**HCHS/SOL**

The baseline examination of HCHS/SOL was carried out as a collaborative study supported by contracts from the National Heart, Lung, and Blood Institute (NHLBI) to the University of North Carolina (N01-HC65233), University of Miami (N01-HC65234), Albert Einstein College of Medicine (N01-HC65235), Northwestern University (N01-HC65236) and San Diego State University (N01-HC65237). The following Institutes/Centers/Offices contributed to the first phase of HCHS/SOL through a transfer of funds to the NHLBI: National Institute on Minority Health and Health Disparities, National Institute on Deafness and Other Communication Disorders, National Institute of Dental and Craniofacial Research (NIDCR), National Institute of Diabetes and Digestive and Kidney Diseases, National Institute of Neurological Disorders and Stroke, NIH Institution-Office of Dietary Supplements. The Genetic Analysis Center at University of Washington was supported by NHLBI and NIDCR contracts (HHSN268201300005C AM03 and MOD03). Genotyping efforts were supported by NHLBI HSN 26220/20054C, NCATS CTSI grant UL1TR000124, and NIDDK Diabetes Research Center (DRC) grant DK063491.

**Health 2000**

The Health 2000 Survey was funded by the National Institute for Health and Welfare (THL), the Finnish Centre for Pensions (ETK), the Social Insurance Institution of Finland (KELA), the Local Government Pensions Institution (KEVA) and other organizations listed on the website of the survey (https://thl.fi/fi/tutkimus-ja-kehittaminen/tutkimukset-ja-hankkeet/terveys-2000-2011/yhteistyokumppanit).

**INGI-CARL/INGI-FVG**

MH 5X1000 funding to IRCCS Burlo Garofolo

**JHS**

The JHS is supported by contracts HHSN268201300046C, HHSN268201300047C, HHSN268201300048C, HHSN268201300049C, HHSN268201300050C from the National Heart, Lung, and Blood Institute and the National Institute on Minority Health and Health Disparities. Dr. Wilson is supported by U54GM115428 from the National Institute of General Medical Sciences.

**Lifelines**

The LifeLines Cohort Study, and generation and management of GWAS genotype data for the LifeLines Cohort Study is supported by the Netherlands Organization of Scientific Research

NWO (grant 175.010.2007.006), the Economic Structure Enhancing Fund (FES) of the Dutch government, the Ministry of Economic Affairs, the Ministry of Education, Culture and Science, the Ministry for Health, Welfare and Sports, the Northern Netherlands Collaboration of Provinces (SNN), the Province of Groningen, University Medical Center Groningen, the University of Groningen, Dutch Kidney Foundation and Dutch Diabetes Research Foundation.

**MESA**

MESA and the MESA SHARe project are conducted and supported by the National Heart, Lung, and Blood Institute (NHLBI) in collaboration with MESA investigators. Support for MESA is provided by contracts HHSN268201500003I, N01-HC-95159, N01-HC-95160, N01-HC-95161, N01-HC-95162, N01-HC-95163, N01-HC-95164, N01-HC-95165, N01-HC-95166, N01-HC-95167, N01-HC-95168, N01-HC-95169, UL1-TR-000040, UL1-TR-001079, UL1-TR-001420, UL1-TR-001881, and DK063491. Funding for SHARe genotyping was provided by NHLBI Contract N02-HL-64278. Genotyping was performed at Affymetrix (Santa Clara, California, USA) and the Broad Institute of Harvard and MIT (Boston, Massachusetts, USA) using the Affymetrix Genome-Wide Human SNP Array 6.0.

**NEO**

The genotyping in the NEO study was supported by the Centre National de Génotypage (Paris, France), headed by Jean-Francois Deleuze. The NEO study is supported by the participating Departments, the Division and the Board of Directors of the Leiden University Medical Center, and by the Leiden University, Research Profile Area Vascular and Regenerative Medicine. Dennis Mook-Kanamori is supported by Dutch Science Organization (ZonMW-VENI Grant 916.14.023).

**ORCADES**

The Orkney Complex Disease Study (ORCADES) was supported by the Chief Scientist Office of the Scottish Government (CZB/4/276, CZB/4/710), a Royal Society URF to J.F.W., the MRC Human Genetics Unit quinquennial programme "QTL in Health and Disease", Arthritis Research UK and the European Union framework program 6 EUROSPAN project (contract no. LSHG-CT-2006-018947).

**PIVUS**

PIVUS was supported by Knut and Alice Wallenberg Foundation (Wallenberg Academy Fellow), European Research Council (ERC Starting Grant), Swedish Diabetes Foundation (2013-024), Swedish Research Council (2012-1397, 2012-1727, and 2012-2215), Marianne and Marcus Wallenberg Foundation, County Council of Dalarna, Dalarna University, and Swedish Heart-Lung Foundation (20120197). Genotyping was funded by the Wellcome Trust under awards WT064890 and WT086596. Analysis of genetic data was funded by the Wellcome Trust under awards WT098017 and WT090532.

**PREVEND**

PREVEND genetics is supported by the Dutch Kidney Foundation (Grant E033), the EU project grant GENECURE (FP-6 LSHM CT 2006 037697), the National Institutes of Health (grant 2R01LM010098), The Netherlands organisation for health research and development (NWO-Groot grant 175.010.2007.006, NWO VENI grant 916.761.70, ZonMw grant

90.700.441), and the Dutch Inter University Cardiology Institute Netherlands (ICIN). N Verweij was supported by NWO VENI (016.186.125). We would like to thank the Center for Information Technology of the University of Groningen for their support and for providing access to the Peregrine high-performance computing cluster.

**PROSPER**

The PROSPER study was supported by an investigator initiated grant obtained from Bristol-Myers Squibb. Prof. Dr. J. W. Jukema is an Established Clinical Investigator of the Netherlands Heart Foundation (grant 2001 D 032). Support for genotyping was provided by the seventh framework program of the European commission (grant 223004) and by the Netherlands Genomics Initiative (Netherlands Consortium for Healthy Aging grant 050-060-810).

**RS**

The Rotterdam Study is supported by the Erasmus Medical Center and Erasmus University Rotterdam; the Netherlands Organization for Scientific Research (NWO); the Netherlands Organization for Health Research and Development (ZonMw); the Research Institute for Diseases in the Elderly (RIDE); the Netherlands Heart Foundation; the Ministry of Education, Culture and Science; the Ministry of Health Welfare and Sports; the European Commission; and the Municipality of Rotterdam. Support for genotyping was provided by the Netherlands Organisation of Scientific Research NWO Investments (nr. 175.010.2005.011, 911-03-012), the Research Institute for Diseases in the Elderly (014-93-015; RIDE2), the Netherlands

Genomics Initiative (NGI)/Netherlands Consortium for Healthy Aging (NCHA) project nr. 050- 060-810. Jacqueline Witteman is supported by NWO grant (vici, 918-76-619). Abbas Dehghan is supported by NWO grant (veni, 916.12.154) and the EUR Fellowship. O.H. Franco works in ErasmusAGE, a center for aging research across the life course funded by Nestlé Nutrition (Nestec Ltd.) and Metagenics Inc. Nestlé Nutrition (Nestec Ltd.) and Metagenics Inc. had no role in design and conduct of the study; collection, management, analysis, and interpretation of the data; and preparation, review or approval of the manuscript. The GWA study was funded by the Netherlands Organisation of Scientific Research NWO Investments (nr. 175.010.2005.011, 911-03-012), the Research Institute for Diseases in the Elderly (014-93-015; RIDE2), the Netherlands Genomics Initiative (NGI)/Netherlands Consortium for Healthy Aging (NCHA) project nr. 050-060-810.

**SardiNIA**

This study was funded in part by the National Institutes of Health (National Institute on Aging, National Heart Lung and Blood Institute, and National Human Genome Research Institute). This research was supported by the Intramural Research Program of the NIH, National Institute on Aging, with contracts N01-AG-1-2109 and HHSN271201100005C; by National Human Genome Research Institute grants HG005581, HG005552, HG006513, HG007089, HG007022, and HG007089; by National Heart Lung and Blood Institute grant HL117626; and by Sardinian Autonomous Region (L.R. no. 7/2009) grant cRP3-154.

**SHIP/SHIP-Trend**

SHIP is part of the Community Medicine Research net of the University of Greifswald, Germany, which is funded by the Federal Ministry of Education and Research (grants no. 01ZZ9603, 01ZZ0103, and 01ZZ0403), the Ministry of Cultural Affairs as well as the Social Ministry of the Federal State of Mecklenburg-West Pomerania, and the network ‘Greifswald Approach to Individualized Medicine (GANI_MED)’ funded by the Federal Ministry of Education and Research (grant 03IS2061A). Genome-wide data have been supported by the Federal Ministry of Education and Research (grant no. 03ZIK012) and a joint grant from Siemens Healthcare, Erlangen, Germany and the Federal State of Mecklenburg- West Pomerania.

**TwinsUK**

TwinsUK is funded by the Wellcome Trust, Medical Research Council, European Union, the National Institute for Health Research (NIHR)-funded BioResource, Clinical Research Facility and Biomedical Research Centre based at Guy’s and St Thomas’NHS Foundation Trust in partnership with King’s College London. This work was supported by the British Heart Foundation (grant number PG/12/38/29615).

**UK Biobank**

This work is supported by Medical Research Council grant MR/N025083/1. MO is supported by an IEF 2013 Marie Curie fellowship.

**ULSAM**

ULSAM was supported by Knut and Alice Wallenberg Foundation (Wallenberg Academy Fellow), European Research Council (ERC Starting Grant), Swedish Diabetes Foundation (2013-024), Swedish Research Council (2012-1397, 2012-1727, and 2012-2215), Marianne and Marcus Wallenberg Foundation, County Council of Dalarna, Dalarna University, and Swedish Heart-Lung Foundation (20120197). Genotyping was funded by the Wellcome Trust under awards WT064890 and WT086596. Analysis of genetic data was funded by the Wellcome Trust under awards WT098017 and WT090532.

**WHI**

The WHI program is funded by the National Heart, Lung, and Blood Institute, National Institutes of Health, U.S. Department of Health and Human Services through contracts HHSN268201600018C, HHSN268201600001C, HHSN268201600002C, HHSN268201600003C, and HHSN268201600004C. Within the WHI, GARNET was funded by the National Human Genome Research Institute through U01-HG005152 (Reiner). GECCO was funded by the National Cancer Institute through R01-CA059045 and U01-CA137088 (Peters). HIPFX was funded by the NHLBI through N01-WH74316 (Jackson). MOPMAP was funded by the National Institute of Environmental Health Sciences through R01-ES017794 (Whitsel) and the University of North Carolina Cancer Research Fund. WHIMS was funded by the NHLBI and NIA through N01-WH44221, 271201100004C and 263200700042C (Shumaker).

**YFS**

The Young Finns Study has been financially supported by the Academy of Finland: grants 286284, 134309 (Eye), 126925, 121584, 124282, 129378 (Salve), 117787 (Gendi), and 41071 (Skidi); the Social Insurance Institution of Finland; Competitive State Research Financing of the Expert Responsibility area of Kuopio, Tampere and Turku University Hospitals (grant X51001); Juho Vainio Foundation; Paavo Nurmi Foundation; Finnish Foundation for Cardiovascular Research ; Finnish Cultural Foundation; The Sigrid Juselius Foundation; Tampere Tuberculosis Foundation; Emil Aaltonen Foundation; Yrjö Jahnsson Foundation; Signe and Ane Gyllenberg Foundation; Diabetes Research Foundation of Finnish Diabetes Association; and EU Horizon 2020 (grant 755320 for TAXINOMISIS); European Research Council (grant 742927 for MULTIEPIGEN project); and Tampere University Hospital Supporting Foundation.

https://www.fredhutch.org/en/labs/phs/projects/cancer-prevention/projects/gecco.html.

12. Genome-wide Association Study to Identify Genetic Components of Hip Fracture. BA3. WHI Study Pages. Women's Health Initiative. https://www.whi.org/researchers/data/WHIStudies/StudySites/BA3/pages/home.aspx.
13. Women’s Health Initiative Study Group. WHI Extension Study Protocol. 2012. https://www.whi.org/researchers/studydoc/Consents/Protocol%202010-2015.pdf.
14. National Institutes of Environmental Health Sciences. Modification of PM-mediated Arrhythmogenesis in Populations. http://projectreporter.nih.gov/project_info_description.cfm?aid=7984809&icde=19283008.
15. Shumaker, S.A. *et al.* The Women's Health Initiative Memory Study (WHIMS): a trial of the effect of estrogen therapy in preventing and slowing the progression of dementia. *Control Clin Trials* **19**, 604-21 (1998).
16. Butler, A.M. *et al.* Novel loci associated with PR interval in a genome-wide association study of 10 African American cohorts. *Circ Cardiovasc Genet* **5**, 639-46 (2012).
17. Chambers, J.C. *et al.* Genetic variation in SCN10A influences cardiac conduction. *Nat Genet* **42**, 149-52 (2010).
18. Holm, H. *et al.* Several common variants modulate heart rate, PR interval and QRS duration. *Nat Genet* **42**, 117-22 (2010).
19. Hong, K.W. *et al.* Identification of three novel genetic variations associated with electrocardiographic traits (QRS duration and PR interval) in East Asians. *Hum Mol Genet* **23**, 6659-67 (2014).
20. Pfeufer, A. *et al.* Genome-wide association study of PR interval. *Nat Genet* **42**, 153-9 (2010).
